# Early-life medial pulvinar disruption drives schizophrenia-relevant prefrontal inhibitory and cognitive deficits in primates

**DOI:** 10.64898/2026.07.23.740162

**Authors:** Jack T. Scott, Sara Fin, Adam P Caccavano, Maxwell Walker, Diego Szczupak, Jordan Jontz, Jihane Homman-Ludiye, William C. Kwan, Kevin Marche, Gordon Fishell, Afonso C. Silva, Kenneth A. Pelkey, Chris J. McBain, James A. Bourne

**Affiliations:** Section on Cellular and Cognitive Neurodevelopment, National Institute of Mental Health, National Institutes of Health; Bethesda, MD 20892, USA; Australian Regenerative Medicine Institute, Monash University; Melbourne, Victoria 3800, Australia; Section on Cellular and Synaptic Physiology, Eunice Kennedy Shriver National Institute of Child Health and Human Development, National Institutes of Health; Bethesda, MD 20892, USA; Department of Neurobiology, University of Pittsburgh; Pittsburgh, PA 15260, USA; Stanley Center for Psychiatric Research, Broad Institute of MIT and Harvard, Cambridge, MA 02142, USA

## Abstract

Schizophrenia is thought to arise from disrupted postnatal maturation of prefrontal circuits, but the developmental events linking early vulnerability to adult cortical dysfunction remain unclear. Here, we tested whether the primate medial pulvinar, a higher-order thalamic nucleus interconnected with prefrontal cortex, contributes to prefrontal maturation. Bilateral medial pulvinar lesions in neonatal marmosets altered adolescent prefrontal diffusion trajectories and produced adult working memory deficits, the latter of which did not follow comparable lesions in adulthood. Early-life lesioned animals showed reduced thalamocortical input to layer 3 parvalbumin interneurons, diminished prefrontal gamma power, reduced parvalbumin expression, and immature-like physiology in fast-spiking interneurons. These findings reveal a developmental window in which thalamic input shapes prefrontal inhibitory maturation, suggesting that some forms of cortical dysfunction in psychiatric disease originate not in the cortex itself, but in its thalamic inputs.

## Main Text

Schizophrenia is a disabling psychiatric disorder in which altered perception and thought are accompanied by persistent impairments in working memory and cognitive control. Symptoms typically emerge in adolescence or early adulthood after a prolonged period in which genetic and environmental risk factors shape maturation of the prefrontal cortex (PFC) (*1*, *2*). This late maturing region of the cortex undergoes dynamic structural refinement (*3*, *4*) accompanied by the consolidation of cell populations toward mature functional states (*5*, *6*). In schizophrenia, convergent PFC abnormalities include the loss or maladaptive pruning of excitatory synapses (*7*, *8*) and suppression of inhibitory interneuron maturation (*9–11*), both of which are strongly linked to cognitive impairment (*1*). However, it remains unclear how early developmental vulnerability is converted into adult PFC circuit dysfunction.

The thalamus is an underexplored, yet plausible source of this developmental instruction as it provides dense input to the developing neocortex. Thalamic neurogenesis and early nuclear organization are largely prenatal (*12*, *13*), yet postnatal thalamocortical connectivity remains dynamic: projections are refined gradually, and track the maturation of cortical areas long into adulthood (*14*). In sensory systems, thalamocortical connections are a primary source of activity-dependent plasticity in the cortex, promoting synaptic complexity and functional identity (*15*, *16*). They can also determine the consolidation of behaviors mediated by the connected cortex during development. In primates, for example, early input from the inferior pulvinar nucleus of the thalamus is required for typical maturation of the dorsal visual stream and coordinated forelimb reaching and grasping (*17*). Similar principles are now emerging in cognitive circuits. Namely, adolescent disruption of the mediodorsal thalamus impairs consolidation of cognitive flexibility and social memory that is associated with maturation of the medial frontal cortex (*18–20*).

In primates, the medial pulvinar (PM) is a strong candidate for shaping maturation of the PFC. PM is a greatly expanded higher-order thalamic nucleus with bidirectional connectivity to areas including the prefrontal, inferior temporal, and posterior parietal cortices (*21*, *22*). Although its postnatal developmental role is unknown, PM is among the thalamic nuclei most consistently implicated in anatomical studies of schizophrenia (*23*). Reduced volume, altered morphology, and lower neuronal number concentrated to the PM suggest a developmental vulnerability of this nucleus (*24–26*). These alterations are associated with poorer cognitive outcomes in adolescence (*25*, *26*), and with reduced pulvinar-frontal functional connectivity in adulthood (*27*). These findings motivate a causal test: does early loss of PM output prevent typical PFC maturation and thereby contribute to adult circuit phenotypes relevant to schizophrenia?

### Excitotoxic lesions model early-life PM disruption

In primates, thalamocortical afferents innervate the cortical presubplate by mid-gestation (*28*), before the major postnatal phase of cortical circuit maturation. To model a perinatal deficit in PM output after initial thalamocortical ingrowth but before postnatal refinement, we produced bilateral PM lesions in common marmosets (*Callithrix jacchus*), a nonhuman primate species with a fetal pulvinar neurogenesis timeline comparable to humans (*29*). We then profiled lesion-induced cortical maturation with diffusion MRI across adolescence into adulthood, followed by cognitive testing, electrophysiological recording, and anatomical assessment (Fig. 1A).

**Figure 1.**
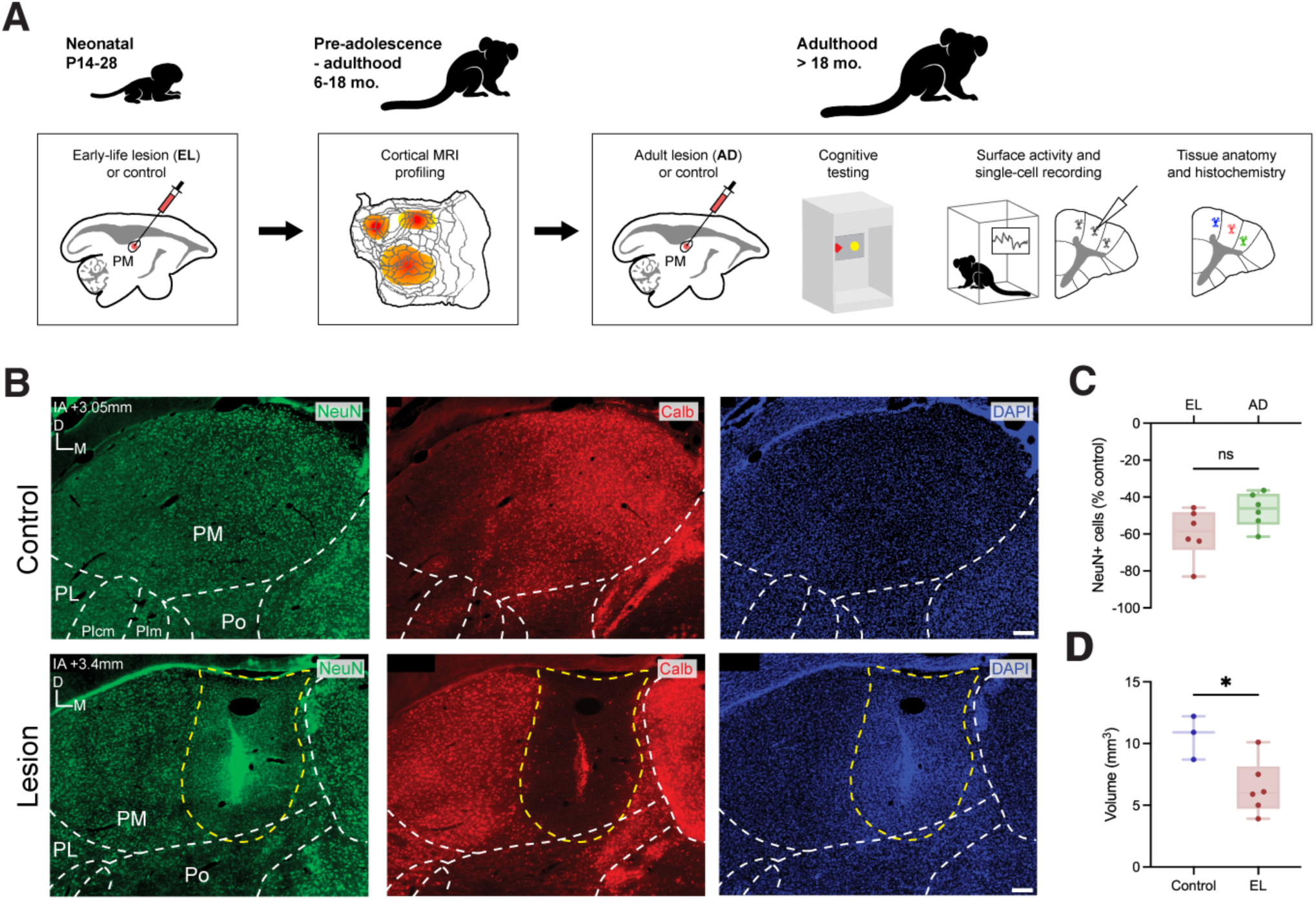
Modeling early PM influences on cortical maturation throughout postnatal development. **A.** Study design used to model early PM maldevelopment and profile its effects on functional cortical maturation throughout adolescence and adulthood, including early-life lesion (EL) and adult lesion (AD) streams. AD animals were not studied in electrophysiology experiments. **B.** Validation of lesions performed in adult animal tissue depicting the reduction in NeuN+ and calbindin+ cells within the lesion core. Lesion images show tissue from an animal in the AD condition. **C.** Quantification of NeuN+ reduction within PM relative to control. **D.** Quantification of PM volume reduction following early lesions. PM: medial pulvinar; PL: lateral pulvinar; PIcm: inferior pulvinar (caudomedial division); PIm: inferior pulvinar (medial division); Po: posterior nuclei. Scale bar = 200 µm. **\*** *p* < 0.05.

Bilateral PM lesions were induced at postnatal day (P)14-28 (*n* = 12; early lesion, EL) by intracerebral injection of excitotoxic NMDA (200 mM, 180 nL). To distinguish developmental mechanisms from general effects of PM injury, identical surgeries were also performed in adult marmosets after 18 months of age (mo.), a time point at which cortical maturation has largely stabilized (*30*) (*n* = 5; adult lesion, AD). Lesion magnitude was quantified posthumously in a subset of adult animals from each cohort (control: *n =* 7 animals, 11 hemispheres; EL: *n =* 3 animals, 6 hemispheres; AD: *n =* 3 animals, 6 hemispheres) using anatomical assessment. Early lesions produced a 59.75% mean reduction in neuronal density based on neuronal nuclear antigen NeuN expression (permutation test, *p* < 0.05) (Fig. 1B, C; Supp. Fig. 1A, F). Lesions appeared particularly biased toward thalamocortical projection neurons, as suggested by a qualitatively extensive loss of calbindin-positive cells within the lesion core (Fig 1B). Because clinical studies of schizophrenia often report reduced PM volume, stereological estimates were used to further confirm a 60.54% mean reduction in PM volume in EL animals relative to control (permutation test, *p* < 0.05) (Fig. 1D). Adult lesions similarly exhibited a 46.98% reduction in neuronal density (permutation test, *p* < 0.001) and did not differ from the EL lesion cohort (permutation test, *p* = 0.078) (Fig. 1B, C; Supp. Fig 1B, F). Glial fibrillary acidic protein (GFAP) immunolabelling further confirmed astrogliosis at the lesion site in both EL and AD cohorts, revealing a glial scar and reactive-like morphology of proximal astrocytes (Supp. Fig 1C-E). Following lesion surgery, animals were differentially assigned to a range of *in vivo* and/or *in vitro* experiments (for a complete summary, refer to Supp. Table 1).

### Early PM connectivity defines the structural maturation of frontal cortex over adolescence

To determine the differential consequences of early PM lesions on postnatal cortical maturation, animals underwent diffusion tensor imaging (DTI) at developmental time points (Fig. 2A). In marmosets, 9 – 18 mo. spans an adolescent sensitive period for neocortical development in which cortical volume declines at the onset of synaptic refinement (*30*) and cell populations acquire mature states (*5*, *6*). DTI scans were therefore performed at pre-adolescent (6 mo.; control: *n* = 6, EL: *n* = 3) and adult (>18 mo.; control: *n* = 9, EL: *n* = 5) time points to characterize changes in cortical microstructure over adolescence. These changes were primarily quantified by comparing fractional anisotropy (FA), a derived measure of voxel-wise diffusion coherence that is sensitive to the cytoarchitectural composition of marmoset cortical gray matter (*31*). Postnatally, cortical gray matter FA follows a U-shaped trajectory, steeply decreasing in childhood and stabilizing in early adulthood during cortical structural reorganization (*32*, *33*). In control animals from 6 mo. to adulthood, statistical surface mapping of voxel-wise differences revealed likewise widespread FA reduction across cortical grey matter, including in both primary and higher areas (voxel-wise *t*-tests, *p* < 0.001; Supp. Fig. 2A). To visualize how this maturation diverged after early lesions, we next compared control and EL animals at both 6 mo. and adulthood. At 6 mo., no differences were reported across the entire cortex (voxel-wise *t*-tests; *p* > 0.001 for all comparisons; Fig. 2B.). By adulthood, cortical gray matter FA had diverged, with higher FA in EL animals localized predominantly to the frontal lobe (voxel-wise *t*-tests; *p* < 0.001; Fig. 2C). FA differences were greatest and most concentrated around areas of the dorsolateral PFC including anatomically defined areas 46D, 46V, 8aD, 8aV, and 8C, with additional effects in ventrolateral PFC areas 47L, 47M and 45, premotor areas 6DR, 6DC, and 6VA, and somatosensory areas 3a and 3ab. Because voxel-wise analyses are sensitive to discrete spatial deviations, we next calculated the mean gray matter FA across the entire PFC and constituent areas for each age and condition (Fig. 2D, E). In controls, FA decreased steeply from 6 mo. to adulthood across nearly all PFC areas (permutation tests; *p* < 0.01 for all significant comparisons) and in the primary motor cortex (M1, area 4ab; permutation test, *p* < 0.001), an adjacent area that has no direct connections to PM (*21*, *22*). In contrast, PFC FA reductions in adolescence after EL were weaker, affecting only M1 and a few PFC areas (9, 46D and 8b; permutation tests, *p* < 0.05). No PFC area differed between control and EL animals at 6 mo. (permutation tests; *p* > 0.05 for all comparisons). In adulthood, however, most PFC areas retained substantially higher FA after EL, including in orbital and medial areas not identified through voxel-based analysis (permutation tests; *p* < 0.05 for all significant comparisons). Together, these findings suggest that early PM lesions suppress typical adolescent microstructural reorganization of the PFC. Developmental reductions in cortical gray matter FA are thought to reflect a shift from predominantly radial organization toward a more complex tangential architecture, including branched basal dendrites and intracortical connectivity (*32*, *34*, *35*). Consistent with this interpretation, the axial-to-radial diffusivity ratio decreased across adolescence in control PFC (permutation tests; *p* < 0.01 for all significant comparisons), indicating a shift toward tangential dominance, but not after EL (permutation tests; *p* > 0.05) (Fig 2F, Supp. Fig 2B-D).

**Figure 2.**
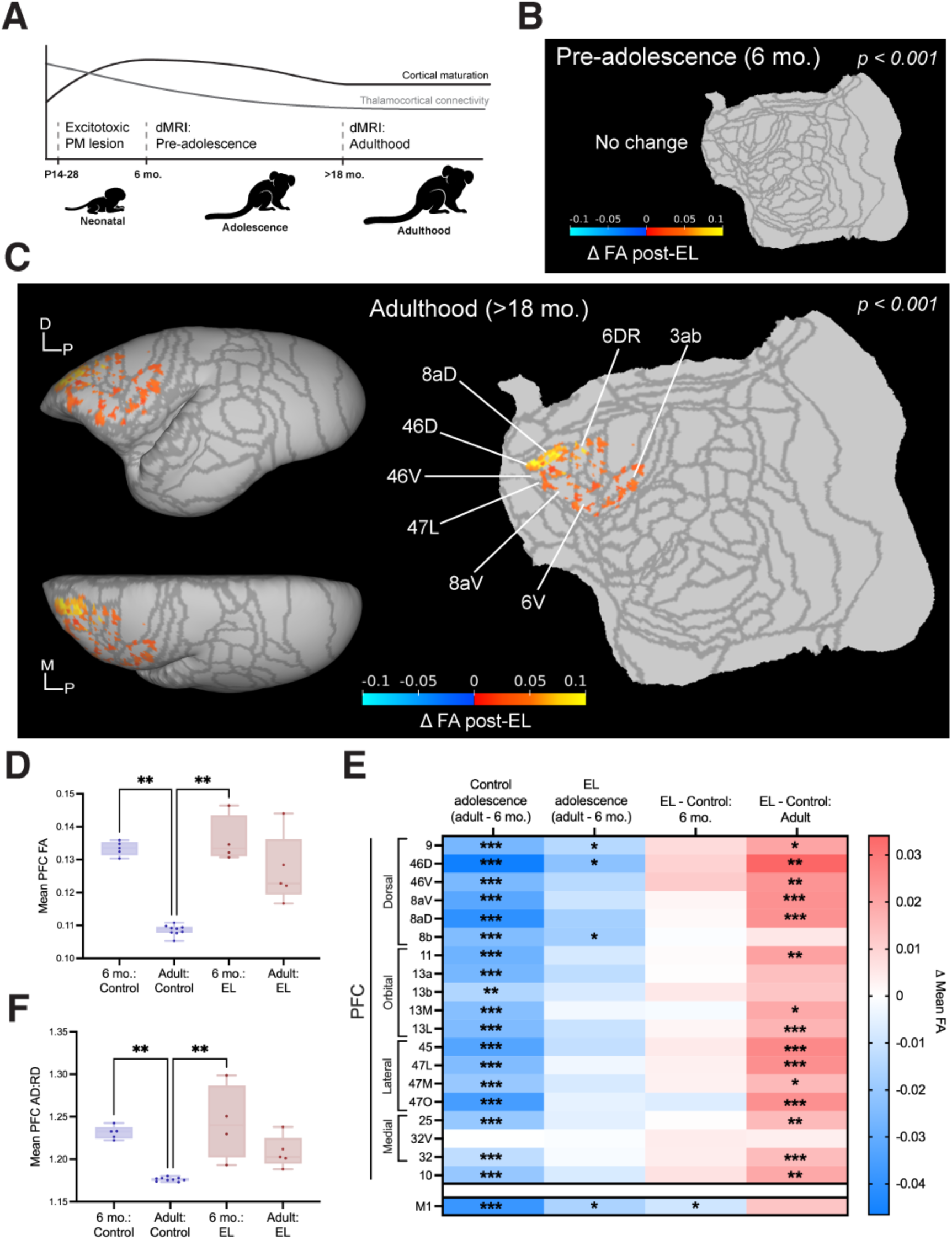
Early-life PM lesions induce divergent microstructural maturation of the frontal cortex over adolescence. **A.** Experimental timeline of diffusion MRI (dMRI) scans. **B.** Statistical flatmap of the neocortical sheet depicting voxel-wise fractional anisotropy (FA) differences in cortical gray matter between EL and controls at 6 months of age (6 mo.). No differences were reported at this pre-adolescent stage. **C.** Statistical flatmap and volume renderings of voxel-wise FA differences in cortical grey matter between EL and controls in adulthood (>18 mo.). Higher values indicate greater FA in EL relative to control. **D, E.** ROI analysis of mean FA within the entire PFC and constituent areas across age and condition. PFC FA reduced over adolescence in controls, but this change was suppressed following early lesions. In contrast, mean FA in M1, an area not recipient of direct PM input, was unchanged in adulthood. **F.** The ratio of axial diffusivity (AD) to radial diffusivity (RD) reduced over adolescence in controls but was unchanged in EL, indicating suppressed tangential dominance of tissue microstructure. **\*** *p* < 0.05, **\*\*** *p* < 0.01, **\*\*\*** *p* < 0.001.

### Postnatal PM connectivity is critical for adult working memory

As schizophrenia-associated PFC maturation is linked to deficits across cognitive control domains, we next asked whether developmental PM disruption produced domain-specific cognitive impairment. Given that regional diffusivity changes were most pronounced in dorsal and lateral PFC and least pronounced in orbital and medial areas, we selected two behavioral tasks that broadly engage the PFC: a delayed match-to-sample task (DMTS) indexing feature-based working memory, and a deterministic reversal learning paradigm indexing cognitive flexibility. Although the PFC performs known roles in working memory, deterministic reversal learning in marmosets is differentially dependent on orbital rather than lateral regions (*36*, *37*). Therefore, we reasoned that these tasks could provide a basic functional interpretation of regional PFC effects in the absence of subcortical contributions. Adult animals performed tasks in *CalliCog,* a homecage-mounted operant chamber that delivers tasks via capacitive touchscreens and rewards desired responses using positive reinforcement (*38*) (Fig. 3A). After automated touchscreen and task training, a subset of animals (control: *n =* 4, EL: *n* = 3; AD: *n =* 3) performed the DMTS. In each trial, animals encoded a ‘sample’ stimulus pseudorandomly sampled from 3 possible geometric shapes, touched the sample twice, and recalled it in the presence of a distractor after a variable delay of 0.5 - 12 s (Fig. 3B). Exponential decay functions were fit to individual animal data to visualize performance as a function of delay (Fig. 3C-D). A generalized linear mixed model (GLMM) was then applied to trial-level data to estimate the fixed effects of the log-transformed delay and lesion condition, while accounting for repeated measures across animals and sessions (Fig. 3E). Increasing delay predicted poorer performance regardless of condition (GLMM with Wald’s test, β = -0.314, *z* = -12.15, 95% CI [-0.362, - 0.263], *p* < 0.0001). Relative to controls, EL animals exhibited significantly poorer overall performance (β = -0.578, *z* = -3.68, 95% CI [-0.886, -0.270], *p* < 0.001), whereas AD did not (β = -0.083, *z* = -0.52, 95% CI [-0.393, 0.228], *p* = 0.601). No significant condition/delay interaction was detected for EL (β = 0.047, *z* = 1.27, 95% CI [-0.025, 0.119], *p* = 0.204) or AD (β = -0.053, *z* = -1.38, 95% CI [-0.129, 0.022], *p* = 0.166), indicating that early lesions caused a generalized working-memory performance deficit rather than a selective increase in delay-dependent decay. Metrics derived from individual decay curves qualitatively supported this interpretation. Relative to controls, EL animals exhibited greater reductions in ‘time to chance’, or the maximum delay at which above-chance performance was achieved (mean difference = 17.10, bootstrapped 95% CI [8.978, 24.181]) compared to AD animals (mean difference = 9.90, bootstrapped 95% CI [1.932, 15.815]) (Fig. 3F); greater reductions in half-life, a readout of decay rate (mean difference = 6.27, bootstrapped 95% CI [2.683, 9.358]), compared to AD animals (mean difference = 3.96, bootstrapped 95% CI [0.598, 6.615]) (Fig. 3G); and a modest reduction in zero-delay performance, an estimate of baseline working memory function independent of delay (mean difference = 7.46, bootstrapped 95% CI [-0.493, 14.050]), compared to AD animals (mean difference = 0.383, 95% bootstrapped CI [-6.774, 8.606]) (Fig 3H.). Thus, early PM lesions impaired working-memory performance in adulthood, with effects consistently larger than those observed after adult lesions. These impairments were notably independent of working memory maintenance, but regardless served to reduce the maximum delay for reliable memory recall.

**Figure 3.**
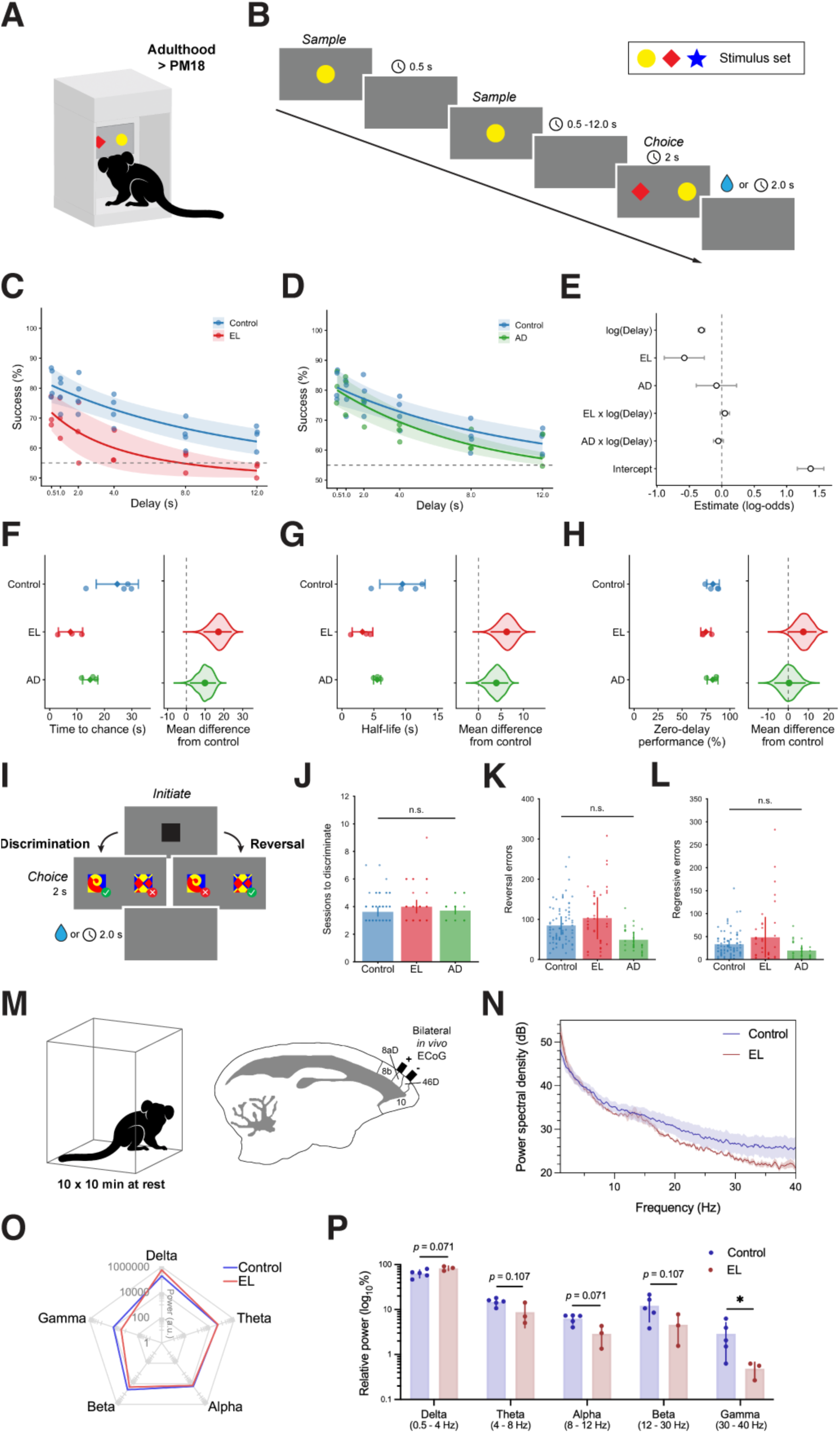
Developmental timing of PM connectivity determines outcomes for adult cognitive control and prefrontal gamma power. **A.** Animals from control, EL and AD cohorts voluntarily performed cognitive tasks in a home cage-mounted platform in adulthood (> postnatal month 18). **B.** Trial representation of the delayed match-to-sample task (DMTS) for feature-based working memory evaluation. Sample stimuli were pseudorandomly selected for each trial from a set of 3 geometric shapes. The delay between the second sample presentation and the choice window ranged from 0.5 to 12 s. **C, D.** Averaged exponential decay functions fitted to the performance of each animal at each delay condition, comparing EL and AD to control. **E.** Regression coefficients of fixed effects from a generalized linear mixed model applied to trial data in the DMTS. Wald Z-tests revealed significant negative predictions for both delay and EL on performance, but not for AD or condition-delay interactions. Data are displayed as estimated coefficients ± 95% CIs. **F-H.** Gardner-Altman estimation plots depicting differences between EL and AD compared to control across metrics derived from exponential working memory functions. Left plots display mean ± SD, right plots display mean difference ± 95% bootstrapped CI and the bootstrapped distribution of the mean difference. **I.** Trial representation of the novel discrimination and reversal learning tasks. Animals discriminated between 10 unique pairs of composite stimuli presented in pseudorandom left or right positions. Completion of each discrimination stage was followed by a reversal in which the stimulus-reward contingencies were deterministically reversed. **J.** Performance in novel discriminations did not differ between conditions. **K-L.** Errors committed following stimulus reversals and errors committed after learning new contingencies did not differ between conditions, despite numerical trends. Data points in the overlaid vertical scatter plot lines represent individual session performance for each animal. **M.** Schematic of experimental design for ECoG recording. EL and control animals were implanted with 2-channel electrodes over the bilateral PFC, and recordings were performed while the animals were awake and at rest for 10 sessions of 10 min each. **N.** Power spectral densities of control and EL recordings computed by multitaper spectral analysis. Data displays average power spectral density +/- SD. **O.** Qualitative shift in distribution of absolute power across delta, theta, alpha, beta and gamma bands in EL compared to control. **P.** Quantification of normalized bandpower per animal comparing control and EL. A significant reduction in gamma (30 – 40 Hz) power was reported in EL relative to control.

To determine whether this impairment generalized to other cognitive processes, a subset of animals (control: *n =* 8, EL: *n* = 4; AD: *n =* 3) underwent novel discrimination and reversal learning tasks to explore feature-based learning and cognitive flexibility. In each task, animals initiated a trial and were presented with pairs of composite stimuli in pseudorandomly selected left and right positions (Fig. 3I). One stimulus was deterministically associated with reward and the other with penalty. During novel discrimination, animals learned to discriminate the rewarded stimulus until reaching proficiency. Across 10 sessions with unique stimulus pairs, conditions did not differ in the number of sessions required to learn initial stimulus-reward contingencies (Kruskal-Wallis test; *H*(2) = 2.053, p = 0.379) (Fig. 3J). Reversal learning was then introduced by swapping stimulus-reward contingencies. After reversal, EL animals committed numerically more errors prior to criterion, but this effect did not reach statistical significance (Kruskal-Wallis test; *H*(2) = 4.762, p = 0.092), whereas AD animals trended in the opposite direction (Fig. 3K). Learning-curve analysis (*39*) categorized trials into three stages of learning: perseverative, learning, and achieving stages (see Methods). Trial proportions across these stages did not differ between conditions (Kruskal-Wallis test; *p* > 0.05 for all tests), though EL animals showed a nonsignificant increase in trials during the achieving phase (Supp. Fig. 3A-C). Consistent with this, regressive errors committed during above-chance performance were not significantly different among groups (Kruskal-Wallis test; *H*(2) = 1.688, *p* = 0.430) (Fig 3L), and errors committed below chance remained comparable to control (Kruskal-Wallis test; *H*(2) = 4.354, *p* = 0.113) (Supp. Fig 3D). Finally, conditions did not differ in the total proportion of win-stay (Kruskal-Wallis test; *H*(2) = 2.117, *p* = 0.347) or lose-shift strategies (Kruskal-Wallis test; *H*(2) = 3.919, *p* = 0.141) after reversal (Supp. Fig 3E). These results indicate that early life PM lesions impaired working memory while leaving deterministic discrimination learning and reversal learning comparatively intact, revealing domain-specific consequences consistent with the observed regional pattern of adolescent prefrontal disruption.

### Adult PFC exhibits reduced gamma power at rest following early-life PM lesions

As behavioral and diffusion MRI findings implicated persistent prefrontal dysfunction after early PM lesions, we next asked whether this phenotype was evident in spontaneous cortical activity. Gamma-band abnormalities are a characteristic physiological feature of PFC dysfunction in schizophrenia and have been linked to cognitive impairment, particularly working memory deficits (*40*, *41*). A subset of adult animals (control: *n =* 5, EL: *n* = 3) was implanted with two-channel electrocorticography (ECoG) electrodes to record bilateral PFC surface activity, with electrodes spanning areas 9, 46D, 46V, 8b, and 8aD. After habituation to the testing chamber, animals underwent 10 recording sessions under controlled conditions while awake and unrestrained (Fig. 3M). Data were processed offline, power spectral densities were estimated using a multitaper method, and absolute band power was calculated using the area under the spectral curve. Inspection of absolute power suggested a shift in EL animals from high to low-frequency power relative to controls, emphasized by dominant shifts in delta (0.5 – 4 Hz) and gamma (30 – 40 Hz), respectively (Fig. 3N-O). After normalization of band power to total spectral power per animal, EL animals revealed a mean reduction in relative gamma power of 83% compared to controls (permutation test, *p* < 0.05), whereas relative power in other bands did not differ between groups (permutation tests, *p* > 0.05 for all comparisons) (Fig. 3P).

### Early-life PM lesions disrupt maturation of parvalbumin-expressing PFC neurons

As early PM lesions altered adult PFC structure, working memory, and resting-state gamma power, we next asked whether these physiological changes were accompanied by abnormalities in cortical parvalbumin (PV)-expressing interneurons. PV is a calcium-binding protein expressed by a major population of fast-spiking cortical interneurons that supports gamma-band synchronization through interactions with pyramidal cells (*42*) and contributes to temporally precise modulation of cognitive circuits (*43*, *44*). We focused on PV interneurons because they receive direct input from thalamic neurons projecting to PFC (*45*, *46*), mature late in postnatal development in primate PFC (*6*, *47*, *48*), and are shaped by activity-dependent mechanisms (*49*, *50*). PV expression was examined in adult control, EL, and AD animals (control: *n =* 6, EL: *n* = 3; AD: *n =* 3) across layer 3 and layer 5/6 of PFC, corresponding broadly to the principal PM-recipient layer and corticothalamic output-associated layers, respectively (*51*) (Figure 4A).

**Figure 4.**
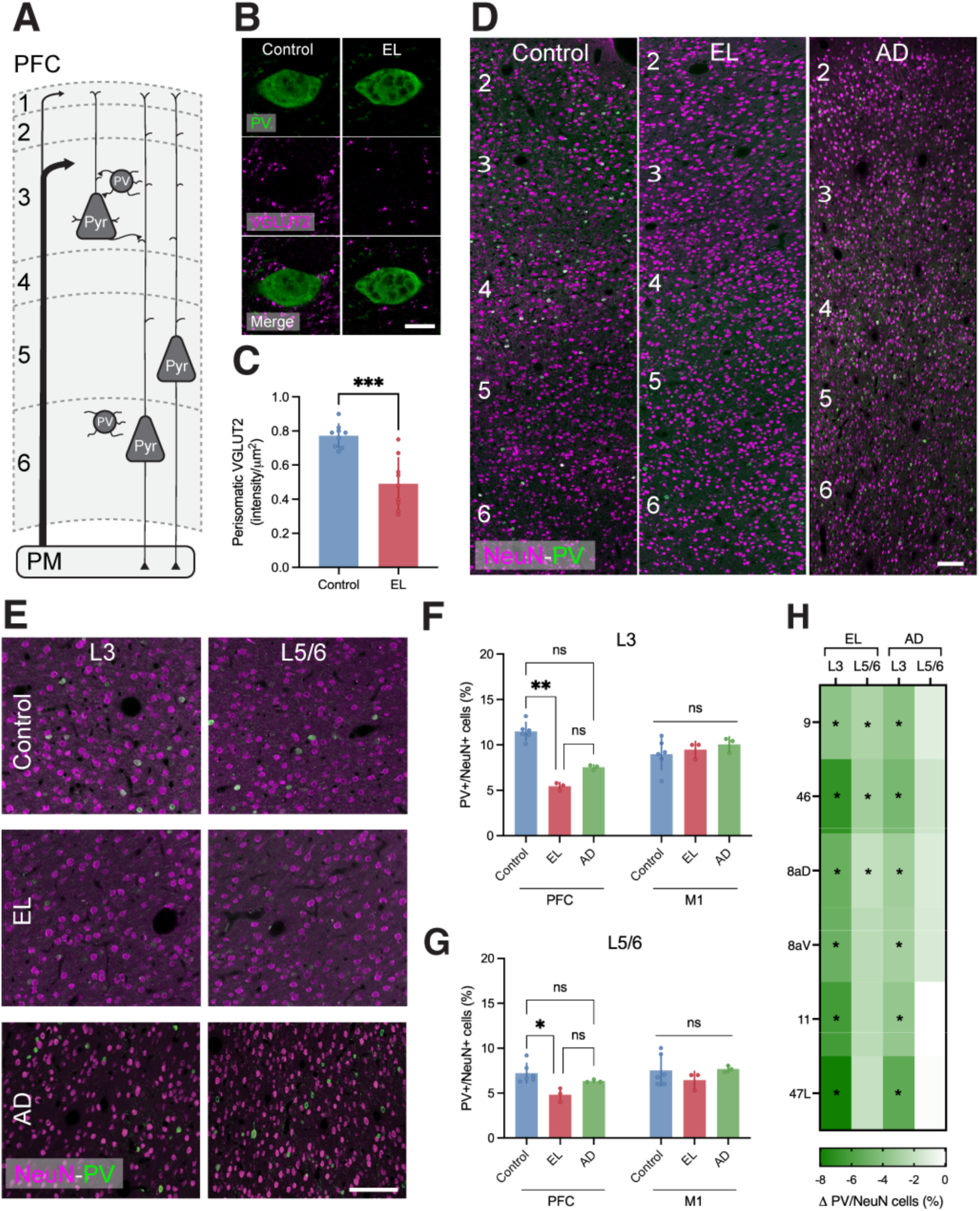
Reduced thalamocortical input to PV interneurons in adult PFC coincides with suppressed PV protein expression. **A.** A representative schematic of wiring between PM and PFC. PM afferents primarily terminate in cortical layer 3, putatively targeting both pyramidal cells and PV interneurons. PV interneurons in layers 5/6 are implicated in the corticothalamic circuit. **B, C.** Reduction of VGLUT2 intensity in the perisomal region of PV+ cells in layer 3 of PV following early lesions, suggesting reduced thalamocortical input. Scale bar = 10 µm. **D-E.** Laminar expression of NeuN and PV immunolabelling in PFC (area 47L) and 10x and 20x objectives. Scale bar = 100 µm. **F, G.** Quantification of the proportion of NeuN+ cells expressing PV in the PFC of both layer 3 and 5/6. The observed changes between conditions were not reported in M1, which lacks direct connectivity with PM. **H.** Cortical area-wise differences in the proportion of PV+/NeuN+ cells compared to control in EL and AD conditions. The areas selected for sampling were chosen to maximize spatial coverage of the PFC. **\*** *p* < 0.05, \***\*** *p* < 0.01, \*\***\*** *p* < 0.001.

To quantify thalamocortical input to PV interneurons, we first measured perisomatic VGLUT2 immunofluorescence around layer 3 PV+ cells. EL animals showed reduced VGLUT2 intensity around PV+ somata in PFC layer 3, consistent with reduced thalamocortical input to this population (permutation test, *p* < 0.001) (Fig 4B, C). We next quantified PV expression among NeuN+ cortical neurons. Across the adult PFC, EL animals showed a broad reduction in NeuN+/PV+ cell density, with the greatest effect in layer 3 (Kruskal-Wallis test with Dunn’s multiple comparisons; *H*(2) = 9.346, p < 0.01) and a lesser, but still significant, effect in layers 5/6 (Kruskal-Wallis test with Dunn’s multiple comparisons; *H*(2) = 8.115, *p* < 0.05) (Fig. 4D-G; Supp. Fig. 4A-B, I). No comparable differences were observed in the control area M1 in layer 3 (Kruskal-Wallis test; *H*(2) = 1.454, p = 0.523) or layer 5/6 of the lesion cohorts (Kruskal-Wallis test; *H*(2) = 2.905, p = 0.252) (Fig. 4F, G). NeuN+ density was stable across both layers (Supp. Fig 4B), suggesting reduced PV protein expression at mature, detectable levels after early PM lesions rather than neuronal loss. The reduction in PV expression was spatially consistent across the PFC. In EL animals, PV+/NeuN+ density was reduced in layer 3 of areas 9, 46, 8aD, 8aV, 11, and 47L and in layer 5/6 of areas 9, 46, 8aD, and 8aV (permutation tests, *p* < 0.05) (Fig. 4H, Supp. Fig. 4E-H). Following adult PM lesions, the phenotype was weaker. Groupwise comparisons did not detect significant differences in PV+/NeuN+ cell density in either layer 3 (*p* = 0.233) or layer 5/6 (*p* = 0.509) in AD animals relative to controls (Fig. 4D-G; Supp. Fig. 4C-D, J). However, exploratory area-wise comparisons revealed reductions in layer 3 across all PFC areas examined (permutation tests, *p* < 0.05), but no reduction in layers 5/6 (permutation tests, *p* > 0.05) (Fig. 4H, Supp. Fig. 3E-H). Together, these findings suggest that PM input most strongly supports PV expression in the principal thalamocortical recipient layer of PFC neurons, and that PM disruption in early life produces broader and more persistent impairments in PFC PV expression than it does in adulthood. These histological changes provide a candidate cellular substrate for the reduced PFC gamma power observed after early lesions and motivated direct testing of PV interneuron physiology.

### Early PM lesions produce immature-like physiological and morphological features in prefrontal PV-class interneurons

As early PM lesions reduced PV protein expression in adult PFC, we next asked whether the PV interneuron population also exhibited altered functional maturation. To target this population independently of mature PV protein detectability, we used S5E2, a regulatory enhancer associated with the SCN1A/Scn1a locus that selectively labels fast-spiking PV interneurons in mammalian cortex (*52*). As S5E2 acts independently of PV gene transcription, we reasoned that S5E2 would label all PV-class neurons including both PV-expressing interneurons and putative PV interneurons in which PV protein had failed to reach mature, immunohistochemically detectable levels. Adult control and EL animals (control: *n =* 2, EL: *n* = 2) were injected in the dorsal PFC with AAV-PHP.eB carrying pAAV(PHPeB)-S5E2-dTom-nlsdTom and processed for slice electrophysiology after viral expression. Single-cell recordings were performed on S5E2+ cells (control: *n* = 51, EL: *n* = 49), and biocytin filled into the cytosol to enable labelling of cell morphology (Fig. 5A). Immunohistochemical analysis of the injection core revealed reductions in PV+ cell density and PV+/SE52+ colocalization between EL and control (permutation test, *p* < 0.05) but no change in S5E2+ cell density (permutation test, *p* = 0.703), suggesting targeting of putative PV+ interneurons in the EL condition (Fig. 5B-C). Recorded S5E2+ cells showed fast-spiking properties characteristic of PV interneurons, exhibiting high sustained firing frequencies > 200 Hz with narrow < 0.5 ms spike half-widths (Fig 5D-E). However, analysis of electrophysiological properties revealed broad differences between EL and control S5E2+ cells (Fig 5D-Fviii, Supp. Table 2). EL cells exhibited a wider action potential (permutation test, *p* < 0.01) (Fig. 5Fi), attributed largely to a slower maximum downstroke (permutation test, *p* < 0.05) (Fig. 5E, Fii) and no difference in maximum upstroke (permutation test, *p* = 0.128) (Fig. 5E, Fiii). In addition, EL cells exhibited an increased time constant (permutation test, *p* < 0.01) (Fig. 5Fiv), increased input resistance (permutation test, *p* < 0.0001) (Fig. 5Fv), and reduced maximum firing frequency (permutation test, *p* < 0.05) (Fig. 5D, Fvi). Consistent with this, EL cells exhibited a reduced capacity to sustain high-frequency firing at higher amplitude current injections (Fig. 5Fvii) and reached depolarization block at lower thresholds relative to controls (Fig. 5Fviii), indicating a failure to maintain firing under strong depolarizing stimulation. In contrast, characterization of spontaneous excitatory post-synaptic currents (sEPSCs) showed no differences in frequency nor amplitude (permutation test, *p* > 0.05 for both comparisons) (Supp. Fig. 5), suggesting that alterations in cell function were not attributable to spontaneous excitatory input. Morphological reconstruction of biocytin-filled S5E2+ cells (control: *n* = 37, lesion: *n* = 31) (Fig. 5G) revealed reduced soma volume (permutation test, *p* < 0.01) (Fig. 5H) and fewer primary neurites (permutation test, *p* < 0.05) (Fig. 5H) in EL cells compared to controls. Together, the preserved density of S5E2+ cells, reduced PV/S5E2 colocalization, altered intrinsic excitability, impaired high-frequency firing, and simplified morphology support the interpretation that early PM disruption leaves adult PFC PV-class interneurons in an immature-like physiological and anatomical state.

**Figure 5.**
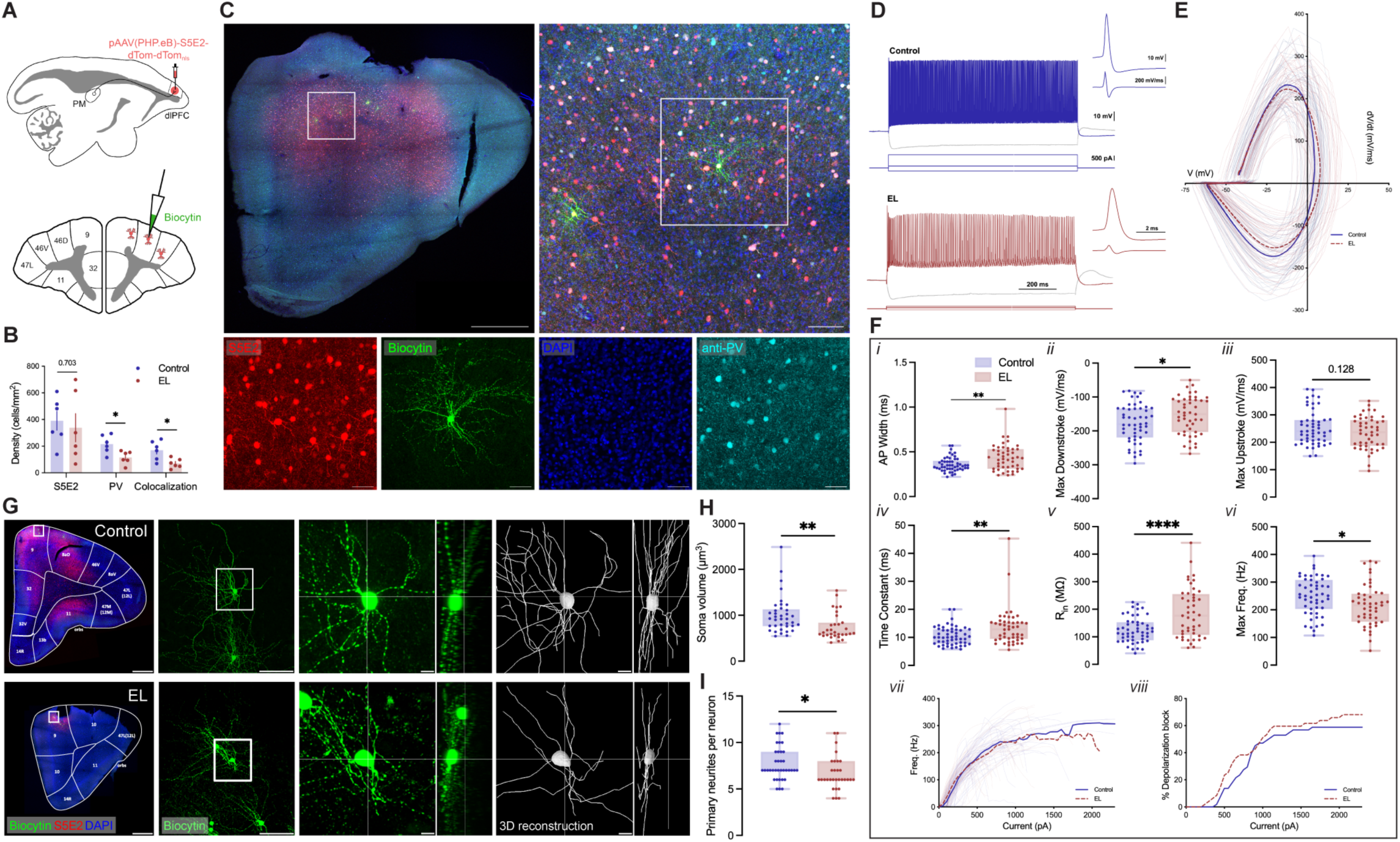
Disrupted maturation of prefrontal PV interneurons following early life PM lesions. **A.** Schematic of the targeting strategy for PV-class interneurons. Both control and EL animals were injected with AAV-PHP.eB carrying a reporter for the S5E2 enhancer selective for PV+ cells. These cells were subsequently identified in slice preparations for patch-clamp electrophysiology and filled with biocytin during recordings to label detailed cell morphology. **B.** Cell counts of immunohistochemical labeling of the injection core, revealing preserved cell density but reduced PV+ and S5E2+/PV+ cell density. **C.** Representative images of targeted cells co-expressing S5E2, PV, and biocytin. Scale bar = 1 mm, 100 µm, 50 µm. **D.** Exemplary spike traces and action potentials of S5E2+ cells in control and EL conditions. **E.** Phase plot visualizing the action potential dynamics of recorded cells. Faint lines depict individual cell action potential dV/dt vs. V, while bold lines show group averages for each condition. **Fi-vi.** Intrinsic electrophysiological properties of recorded cells. **Fvii, viii.** Current-evoked firing properties depicting the firing frequency and depolarization block between control and EL conditions. **G.** 3-dimensional reconstruction of biocytin-labelled cells. Scale bar = 1 mm, 100 µm, 10 µm. **H, I.** Quantification of cell morphology characteristics in control and EL conditions. **\*** *p* < 0.05, \***\*** *p* < 0.01, \*\***\*\*** *p* < 0.0001.

## Discussion

In this study, we used a cell-to-system strategy to test whether PM input is required for typical postnatal maturation of the primate PFC. Early-life PM lesions disrupted the typical organization of frontal cortical gray matter microstructure and produced an adult phenotype of impaired prefrontal inhibition, characterized by reduced thalamocortical input to layer 3 PV+ cells, reduced neuronal PV protein expression, and immature-like intrinsic and morphological properties in S5E2-targeted PV-class interneurons. At the system and behavioral levels, early lesions reduced awake PFC gamma power and impaired feature-based working memory while comparatively sparing discrimination learning and cognitive flexibility. These effects were weaker or absent where adult lesion comparisons were applied, supporting a dependence on developmental timing. Together, these convergent anatomical, physiological, and behavioral findings support a model in which early higher-order thalamic input instructs the maturation of primate PFC inhibitory circuits needed for mature cognition. Outcomes for PV interneuron phenotype, gamma-band activity, and working memory closely align with schizophrenia-relevant PFC dysfunction, identifying PM–PFC development as a candidate mechanism linking early thalamic vulnerability to adult cognitive impairment.

### Implications for prefrontal pathology in schizophrenia

Consistent hallmarks of schizophrenia include PFC synaptic and inhibitory abnormalities (*7*, *9–11*), thalamocortical dysconnectivity (*27*, *53*, *54*), and broad cognitive impairment. While the cause of cognitive impairment remains poorly defined, genetic and environmental factors converge on a perinatal critical period that confers vulnerability on the developing PFC (*55*, *56*). The PM has been proposed as a candidate locus for this early developmental vulnerability (*51*, *57*), and postmortem and neuroimaging studies implicate pulvinar/PM structure in schizophrenia and related psychosis (*23*, *25–27*, *58*). In this study, we provide the first causal evidence that an early PM deficit reproduces features that align with schizophrenia-associated pathology: impaired working memory (*59*), altered prefrontal gamma activity (*60*, *61*), and reduced cortical PV expression (*10*). In addition, PV-class interneurons in adult animals with early PM lesions approximated an immature functional state presumed to occur in the disorder but never demonstrated clinically. For example, PV-class cells in early-lesioned animals exhibited reduced branching and neurite length; greater input resistance, time constant and action potential duration; and reduced maximum firing frequency; all of which distinguish pre-adolescent PV interneurons from mature ones (*62–64*). Collectively, these data do not model schizophrenia as a syndrome, nor do they imply that PM lesions are a common clinical cause. Rather, they show that early-life disruption of specific higher-order thalamic input implicated in the disorder is sufficient to assemble a schizophrenia-relevant prefrontal hallmarks. This places thalamocortical maturation upstream of the interneuron and oscillatory abnormalities often attributed to local cortical microcircuits (*65*, *66*), and suggests a disease mechanism in which early thalamic vulnerability is amplified into adult cognitive dysfunction.

### A unique role for primate thalamus in cortical maturation

Thalamic nuclei can be broadly classified into two types by their dominant source of presynaptic excitation, or “driver” input (*67*, *68*). First-order nuclei receive their primary driver input from the sensory periphery, whereas higher-order nuclei, such as PM, receive major driver input largely from layer 5 of the cortex. The role of first-order thalamus in sensory cortical development is well established (*15*, *16*, *69*), but relatively little is known about the developmental contribution of higher-order thalamus. Rodent studies of the mediodorsal nucleus (MD), a major source of thalamic input to the frontal cortex, have begun to define such mechanisms. Transient MD inhibition in the mouse during adolescence (P35 – 42) reduced L2/3 pyramidal cell excitability, medial frontal cortical activity, and social memory in adulthood (*19*). More extended MD inhibition (P20 – 34 or P20 – 50) impaired working memory rule acquisition and cognitive flexibility (*18*) and alter excitation/inhibition balance in discrete prefrontal projections from the thalamic input and output layers (*20*). Our findings similarly show poorer adult working memory after early disruption of higher-order thalamic input. However, PM lesions produced a broader deficit in working memory performance, rather than the rule acquisition phenotype reported after MD inactivation (*18*). Outcomes for inhibitory physiology also differed: early PM lesions reduced population-level gamma power and shifted PV-lineage interneuron firing properties, whereas MD inactivation studies reported no change in PV interneuron excitability (*19*) or in task-evoked gamma power associated with PV function (*18*, *50*). These differences suggest that PM, and perhaps the expanded primate pulvinar complex more generally, has a distinct role in shaping cortical inhibition during postnatal development.

### Medial pulvinar function may arise from evolutionary development

In this study, we show that early postnatal PM input is required for typical maturation of inhibitory circuits in the PFC. This raises an important question: why is this process so sensitive to PM disruption despite the presence of other thalamic inputs to PFC? MD has substantially denser connections with PFC than PM (*70*), has itself undergone elaboration in parallel with primate PFC specialization (*71*), and exerts strong control over cortical PV interneurons (*46*, *72*, *73*). We propose that the developmental effect of PM may depend less on the strength of its feedforward projection and more on the timing and origin of the signals it relays during sensitive periods. The primate neocortex comprises many specialized areas that mature according to a heterogeneous postnatal timetable (*3*, *30*, *74*). This heterochronic maturation is linked to dynamic thalamocortical connectivity, which develops sequentially along a sensorimotor-to-association axis (*14*, *75*, *76*). Timing also appears to reflect the functional demands of the developing organism. For example, transient connections through the primate inferior pulvinar provide an early route by which retinal information can shape higher visual areas needed for visuomotor coordination in early life, at a stage when visual information is still sculpting the primary visual cortex (*17*). By analogy, PM similarly receives dynamic putative driver inputs from layer 5 of temporal and parietal association cortex (*57*), positioning it to integrate higher dimensional sensory signals to the maturing PFC. Under this model, early PM lesions do not simply remove an adult thalamic input; they remove a time-locked developmental scaffold that MD and other thalamic pathways cannot fully replace. Thus, PM may facilitate an extended sensitive period for cognitive development by coordinating association cortex drive with the maturation of prefrontal inhibitory circuits.

Here, we identify a developmentally timed higher-order thalamocortical mechanism through which early PM input supports the maturation of primate prefrontal inhibitory circuitry and the refinement of adult cognition. Early-life PM disruption produced a constellation of schizophrenia-relevant prefrontal phenotypes, including altered frontal cortical microstructural maturation, impaired working memory, reduced gamma-band activity, diminished PV expression, and immature-like fast-spiking interneuron physiology. These findings position PM–PFC development as a candidate nexus between early thalamic vulnerability and later cortical dysfunction, and suggest that higher-order thalamic nuclei do more than modulate mature cortical processing: they may provide temporally specific developmental scaffolds through which association cortices acquire mature inhibitory properties for cognition. Defining the genetic, environmental, and circuit-level factors that perturb these thalamocortical programs may reveal mechanisms of cognitive impairment across neurodevelopmental disorders and identify sensitive windows for intervention.

## Materials and Methods

### Subjects

A total of 40 common marmosets (*Callithrix jacchus; n =* 40; 19 female, 21 male) were used in the current study. Biological sex was not a selection criterion for individual experiments, as most experiments depended on the timing of available births. Age was variable and experiment-dependent, ranging from infancy (postnatal day (P)14-28) through pre-adolescence (6 mo.) and adulthood (18-72 mo.). A detailed summary of all study subjects, including sex, cohort, and experimental procedures, is provided in Supplementary Table 1.

Animals were housed between 3 different facilities. In Facility 1, animals (*n* = 20) were housed indoors in vertically stacked double cages joined by a tunnel (200 cm x 78 cm x 78 cm, 13 cm elevation). In Facility 2, animals (*n* = 16) were housed in cages (144 cm x 75 x 64 cm, 40 cm elevation) equipped with access tunnels to a larger outside enclosure, which could be freely explored during daytime hours when not under experimentation. A small subset (*n* = 3) was held at both Facility 1 and 2 across the course of the study. In Facility 3, animals (*n* = 7) were housed in conditions akin to Facility 1. All animals were reared by their parental dyad and were not relocated from their family unit to separate housing until at least 12 mo. Once relocated, animals were either single- or pair-housed and retained visual and audial access to other marmosets in the vivarium. All facilities operated on a 12-hr light/dark cycle (07:00 – 19:00) and maintained controlled ambient conditions (26-28 °C, 40–60% humidity). Animals were fed a rotating diet comprising various fruits, nuts, insects, and formulated primate pellets or chow daily after experimentation, and food was removed from the cage prior to testing. *Ad libitum* access to water was provided at all times. In-cage behavioral testing was performed in Facilities 1 and 2 only between 09:00 and 17:00 daily, Monday to Friday, with each individual animal given access to an operant chamber once daily for at least 2 hours, depending on schedule and engagement.

All animal procedures in Facility 1 were fully compliant with the Guidelines for the Care and Use of Laboratory Animals by the National Institutes of Health and approved by the Animal Care and Use Committee of the National Institute of Mental Health. All animal procedures in Facility 2 were conducted in accordance with the Australian Code of Practice for the Care and Use of Animals for Scientific Purposes and were approved by the Monash University Animal Ethics Committee, which also monitored animal welfare. All experimental procedures in Facility 3 were in accordance with ethical guidelines for animal testing established by the University of Pittsburgh Institutional Animal Care and Use Committee.

### Surgery

#### Intracerebral microinjection

Animals were anesthetized with an intramuscular injection of alfaxalone (6-8 mg/kg; Jurox) and diazepam (1-3 mg/kg; Pfizer) and maintained with inhalant isoflurane (1-2% in oxygen). Atropine (20 μg/kg; Pfizer) and dexamethasone (0.2 mg/kg; Ilium) were also administered intramuscularly to reduce mucosal secretions, control heart rate, and prevent cerebral edema. Bupivacaine (0.1 – 0.3 mL; Aspen) was administered subcutaneously for local anesthesia, and meloxicam (0.1-0.6 mg/kg) subcutaneously for analgesia. Vitals, including heart rate, blood oxygen saturation, and external temperature, were monitored throughout the procedure and kept within optimal ranges.

Animals were then prepared for surgery and secured to a stereotaxic frame. Following a scalp incision, injection trajectories were identified and marked on the cranium. This was achieved via two different methods: either animals were transported for structural MRI scanning mid-surgery and trajectories were manually calculated based on fiducial markers as described previously (*77*); or trajectories were determined based on the three-dimensional alignment of the cranium to pre-acquired structural MRI/CT data using the BrainSight Vet Robot system (Rogue Research Inc.). Burr-hole craniotomies and durotomies were then performed over the marked sites, and a 5 μL microsyringe (Neuros Syringe (Hamilton; Cat# 65460-03) fitted with a customized 33G 15-degree beveled needle (Hamilton; Cat# 65461-02)) was advanced into the parenchyma using a micromanipulator (Kopf Instruments) or robotic arm (Rogue Research Inc.). For excitotoxic lesions of bilateral PM in infant (*n* = 12) and adult (*n* = 5) animals, 180 nL of *N-*methyl-D-aspartate (NMDA; Sigma Aldrich) was injected at a rate of 180 nL/min in each hemisphere. For S5E2 labeling, 500 nL of AAV-PHP.eB carrying pAAV(PHPeB)-S5E2-dTom-nlsdTom (titer = 1 x 10^13^ vg/mL) was injected at a rate of 180 nL/min into both areas 9 and 46D in the left prefrontal cortex (PFC). The needle remained in place for a further 10 min before being slowly retracted to allow the infusion to dissipate. Cranial and skin repair were then performed and animals recovered from anesthesia, followed by a minimum 14-day post-operative recovery period in the home cage.

#### ECoG radiotelemetry implantation

Adult animals in control (*n* = 5), early lesioned (EL; *n* = 3), and adult lesioned (AD; *n* = 4) cohorts were implanted with intracranial electrodes to record 2-channel electrocorticography from the bilateral PFC. Procedures for surgical preparation, anesthesia, vitals monitoring, and postoperative recovery were identical to those of intracerebral injection. For implantation of the radiotelemetry device, a horizontal dermal incision was made between the dorsal scapulae in the back of the animal, and a subcutaneous pocket was created adjacent to the incision. The radiotelemetry transmitter (HD-S02, Data Sciences International; 28 x 17 x 8 mm) was inserted into the pocket, and the attached wires were routed subcutaneously from the implant to a second surgical incision site made along the midline of the scalp. The first incision in the back of the animal was then sutured. Burr hole craniotomies were performed in the cranium, and 2 iridium oxide intracranial screw electrodes were driven in anterior and posterior positions above both the left and right PFC to contact the dura mater. Electrode placement was aligned to stereotaxic coordinates (16.5 mm and 15.5 mm anterior to the interaural line, +/- 2 mm lateral to the midline) and then adjusted slightly for brain morphology so that electrodes spanned cortical areas 9, 46D, 46V, 8b and 8aD, as determined by the digitized atlas of Paxinos et al. (*78*). Wires from the dorsal telemetry device were then anchored to the screws, which were then insulated with dental cement (C&B Metabond, Parkell Inc.). Skin repair was then performed, and animals recovered from anesthesia, followed by a minimum 14-day post-operative recovery period in the home cage.

### MRI and CT

#### Acquisition

MRI scans were acquired using 3 scanner configurations:

1. A 9.4 T horizontal 20 cm bore MRI scanner (Bruker Biospin Corp) equipped with a Bruker BioSpec Avance III HD console and the software package Paravision 6.0.1 (Bruker Biospin Corp) housed at Monash Biomedical Imaging, Monash University. Radiofrequency transmission was accomplished with an 86 mm inner diameter volume coil. A 4-ch phased-array receive coil (RAPID Biomedical) was used for radiofrequency receiving.
2. A 7 T horizontal 30 cm bore MRI scanner (Bruker Biospin Corp) equipped with a Bruker BioSpec Avance Neo console and the software package Paravision 360 (version 3.6; Bruker Biospin Corp), housed at the Neurophysiology Imaging Facility, National Institute of Mental Health. Radiofrequency transmission was accomplished with a 13.4 cm inner diameter volume coil. A custom-built 8-ch phased-array coil was used for radiofrequency receiving.
3. A 9.4 T 30 cm horizontal bore MRI scanner (Bruker BioSpin Corp) equipped with a Bruker BioSpec Avance Neo console and the software package Paravision 360 (version 3.6; Bruker BioSpin Corp), housed in the Department of Neurobiology at the University of Pittsburgh. Radiofrequency transmission was accomplished with a custom 150 mm inner diameter coil. A custom-built 16-channel phased-array coil was used for radiofrequency receiving.

CT scans were acquired using a Vimago HU CT scanner (Epica International). Structural images were obtained for non-quantitative surgical targeting only, and included T2-weighted images from a 2D RARE sequence acquired in the axial plane: TR = 12,000 ms, TE = 48 ms, FOV = 40 x 36 mm^2^, matrix size = 200 x 180, in-plane resolution = 0.2 x 0.2 mm^2^, slice thickness = 0.4 mm, RARE factor = 8, number of averages = 4, scan time = 30 min; or T1-weighted images from a 3D MPRAGE sequence: TR = 6 ms, TE = 2.8 ms, TI = 1.2 ms, FOV = 36 x 36 x 32 mm^3^, matrix size = 144 x 144 x 128, resolution = 0.25 mm isotropic, number of averages = 1, scan time = 10 min. CT images were obtained with an isotropic voxel size of 150 μm. Diffusion-weighted images were obtained using the following sequences for each of the 3 MRI scanner configurations:

1. a 3D 4-shot spin-echo EPI sequence, TR = 300 ms, TE = 32 ms, FOV = 36 x 28 x 18 mm^3^, matrix size = 72 x 56 x 36, voxel size = 0.5 x 0.5 x 0.5 mm, number of averages = 2, 2 *b* = 0 and 64 *b* = 4000 s/mm^2^ were each collected twice with reversed phase-encoding directions (superior-inferior and inferior-superior), scan time = 95 min.
2. a 2D single-shot spin-echo EPI sequence, TR = 5100 ms, TE = 39 ms, FOV = 28 x 36 x 19 mm^3^, matrix size = 56 x 72 x 38, slice thickness = 0.5 mm, 38 slices, number of averages = 1, GRAPPA acceleration factor of 2, 8 *b* = 0, 64 *b* = 1000, and 128 *b =* 2000 s/mm^2^ were each collected twice with reversed phase-encoding directions (right-left and left-right), scan time = 17 min.
3. a 2D single-shot spin-echo EPI sequence, TR = 5100 ms, TE = 38 ms, FOV = 42 × 35 mm, matrix size = 84 × 70, slice thickness = 0.5 mm, 50 slices, number of averages = 1, GRAPPA acceleration factor of 2, 8 *b* =0, 64 b = 1000 s/mm^2^, and 128 *b* = 2000 s/mm^2^ were each collected twice with reversed phase-encoding directions (right-left and left-right), scan time = 20 min

#### Preprocessing

Diffusion-weighted images were denoised using the *dwidenoise* tool from MRtrix 3.0 (*79*) and corrected for susceptibility- and eddy-current-induced distortions using *topup* and *eddy* from FSL 6.0 (*80*). A diffusion tensor model was fit to the datasets using the *dtifit* tool from FSL. Resultant fractional anisotropy (FA) maps were nonlinearly registered using ANTs (*81*) to a template space from the Marmoset Brain Mapping project (version 2.0) (*82*) and down-sampled to 0.5 mm isotropic resolution, where necessary. To harmonize the data across scanner configurations, control data from configurations 2) and 3) were normalized to age-matched data from configuration 1) using the neuroHarmonize package (*83*). Data from lesioned animals obtained using configuration 2) were transformed by applying the model trained on control animals from the same configuration.

### Behavior

#### Touchscreen training

Touchscreen training was administered using an automated, home cage-integrated operant chamber (*CalliCog*) following a protocol detailed elsewhere (*38*). In brief, animals performed a series of 7 task phases, beginning with responses to a large colored stimulus (RGB red, blue, or yellow) that was progressively reduced in size and later varied in position.

Per trial, correct responses delivered a liquid reward (marshmallow solution, 20% w/v in water, ∼150 µL, paired with an 800 Hz tone), and incorrect responses initiated a penalty timeout (2 s, 100 Hz tone). Progression from each phase occurred once animals successfully completed three consecutive sessions of 50 trials with an 80% success rate.

#### Novel discrimination/reversal learning task

Animals were progressed to a novel discrimination and reversal learning task after completing touchscreen training. As described previously (*38*), the task involved individual sessions that each included a novel discrimination and reversal learning stage. During novel discrimination, animals responded to a stimulus to initiate a trial (a black square) and were presented with a pair of novel stimuli presented pseudorandomly in left and right positions. Stimulus pairs consisted of 250 x 250 px geometric images on a white background and were deterministically associated with reward and penalty (2 s timeout). A response to either stimulus concluded the trial; no response within 2 s terminated the trial. Animals performed this task until proficiently discriminating the correct stimulus, as determined by a ≥80% success rate over 3 consecutive sessions of 50 trials. Upon achieving this criterion, the reversal learning phase of the task was initiated, and stimulus-reward contingencies were immediately reversed. Animals performed the task under the new contingencies until achieving 90% success over a 100-trial rolling window. In total, animals were presented with 3 pairs of novel stimuli for initial discrimination training, followed by 10 pairs for the novel discrimination/reversal learning task.

#### Delayed match-to-sample task

Animals were trained on the delayed match-to-sample task (DMTS) following completion of reversal learning experiments. First, animals progressed through 4 individual training phases, as described previously (*38*). In brief, training involved the central presentation of a sample stimulus to be recalled, pseudorandomly assigned as either a yellow circle or a red diamond, with an invisible 250 x 250 px hit box. Animals responded to the sample, and following a 0.5 s delay, were presented with a forced choice window containing the sample and a distractor (the remaining unassigned stimulus) in pseudorandom left and right positions. The trial concluded upon selecting either stimulus, resulting in a reward or a 3-second penalty timeout. No response within 3 s after the forced-choice window terminated the trial. During training, the following parameters were progressively added: 1) the sample stimulus was presented twice to ensure visual attention to features; 2) a third, central position was introduced as a possible location for a stimulus in the forced choice window; 3) a third stimulus type, a blue star, was added to the pool of stimuli to be assigned a sample or a distractor. Animals were considered to have completed training once they performed this version of the task to ≥80% success over 3 consecutive sessions of 50 trials, and were subsequently progressed to the testing phase of the task. In the testing phase, a variable delay period was added between the final presentation of the sample and the forced choice window. Initially, shorter delays of 0.5, 1, 2, and 4 s were pseudorandomly interleaved, followed by progressively longer periods of 8 and 12 s as animals engaged over more trials. In total, animals performed 400 trials each with variable delays of 0.5 – 4 s, 300 trials with a variable delay of 8 s, and 150 trials with a variable delay of 12 s.

### ECoG

#### Awake recordings

Animals underwent daily training for 30 min to habituate to a novel testing environment prior to electrocorticography (ECoG) implantation. First, animals were permitted to freely access a transparent acrylic transport chamber mounted on the home cage. Next, the door to the chamber was gradually closed with the animal inside, and food rewards (marshmallows) were presented through reward ports to habituate animals to the chamber. Upon consistently presenting behavioral signs indicative of normal exploration and in the absence of indicators of stress (e.g. piloerection, bobbing, rapid movements, vocalizations), animals were transported within the chamber to an external soundproofed testing room and placed in an opaque container. Here, animals were habituated to the altered environment using positive reinforcement until they exhibited no stress-like behaviors lasting >15 s over a total of 15 min, without food rewards. After recovery from surgical implantation, animals were retrained in this procedure until they met this criterion. Recordings were performed in this testing environment in the absence of a human investigator across 10 consecutive daily 10-min sessions. Critically, recordings were obtained wirelessly from a receiver underneath the transport box, and the implanted radiotelemetry device was magnetically activated without requiring direct handling. Animals were remotely monitored by camera throughout recordings, and sessions were terminated and considered null if stress-like behaviors were observed and lasted >15 s.

#### Preprocessing

2-channel recordings corresponding to left and right PFC activity were acquired during sessions at a sampling rate of 500 Hz and visualized using Ponemah software (version 6.5.1, Data Sciences International). The raw data were then exported and processed offline using custom Python code. Firstly, segments of signal dropout were detected when animals moved beyond the telemetry receiver, and these were removed from the data. Next, transient increases in broadband power associated with temporalis muscle contractions were identified and removed from the data, as described previously (*38*). The valid data were then fed through a notch filter (50 or 60 (+/- 5) Hz, depending on the recording location) to remove line noise contaminant, and bandpass filtered from 0.5 – 100 Hz with a zero-phase, sixteenth-order Butterworth filter. All data used for subsequent analysis was the average signal from both channels.

### Perfusion and tissue processing

Animals were overdosed with sodium pentobarbital (100 mg/kg, intraperitoneal), and upon loss of corneal reflex and cardiac arrest were transcardially perfused. Tissue assigned to patch clamp electrophysiology was perfused with ice-cold, sucrose-substituted artificial cerebrospinal fluid (SSaCSF) containing (in mM): 90 sucrose, 80 NaCl, 3.5 KCl, 1.25 NaH_2_PO_4_, 24 NaHCO_3_, 10 glucose, 0.5 CaCl_2_, and 4.5 MgCl_2_ saturated with carbogen (95% O_2_, 5% CO_2_), with osmolarity 310-320 Osm. Tissue was blocked and coronally sectioned (300 µm) on a VT-1200S vibratome (Leica Microsystems), then transferred to a submerged incubation chamber containing oxygenated warmed (32-34°C) SSaCSF for 30 minutes, then maintained at room temperature for up to 72 hours. For all other experiments, tissues were perfused with ambient-temperature heparinized isotonic saline (0.9% w/v), followed by paraformaldehyde (4% w/v). Brains were dehydrated in increasing concentrations of sucrose (10%, 20% and 30%) over 3 days, flash-frozen over liquid isopentane, and stored at -80 °C. A cryostat (Leica CM3050 S) was then used to section tissue to 40 μm coronal sections in 4 series, and sections were stored in cryoprotectant at -20 °C.

### Patch clamp electrophysiology

Acute slices were transferred to an upright microscope (Olympus BX51Wl) and perfused with oxygenated extracellular SSaCSF containing (in mM): 130 NaCl, 3.5 KCl, 1.25 NaH_2_PO_4_, 24 NaHCO_3_, 10 glucose, 2.5 CaCl, 1.5 MgCl_2_, with osmolarity 300-310 Osm, with a flow rate of 2-3 mL/min and a temperature of 30-33 °C. Individual cells were visualized with a 40x objective using red fluorescence and IR-DIC microscopy. Electrodes were pulled from borosilicate glass (World Precision Instruments) to a resistance of 3-5 MΩ with a vertical pipette puller (Narishige PC-10). Whole-cell patch-clamp recordings were made with a Multiclamp 700B amplifier (Molecular Devices), with signals digitized at 20 kHz (Digidata 1440A, filtered at 3 kHz). Recordings were made using a Windows computer with pClamp 10.7 (Molecular Devices). In voltage-clamp recordings, uncompensated access resistance (R_A_) was monitored consistently with 5 mV voltage steps. High and unstable R_A_ recordings were discarded, as were recordings with unstable leak current.

Intrinsic membrane and firing properties were recorded in fluorescently tagged cells with a standard K-Gluconate internal solution containing (in mM): 150 K gluconate, 0.5 EGTA, 3 MgCl_2_, 10 HEPES, 2 ATP•Mg, 0.3 GTP•Na_2_, and 0.3% biocytin (pH corrected to 7.2 with KOH, 285-290 Osm). Resting membrane potential (RMP) was measured in a tight cell-attached configuration voltage-clamped to +60 mV during 100 ms voltage ramps (from +100 to -200 mV) every 5 s, followed by a whole-cell current-clamp configuration with holding current I = 0 after break-in. Membrane time constant (tau) was measured by 20 repeated 400 ms hyperpolarizing current pulses of -20 pA. Input resistance (R_in_) was determined by recording the response to 20 increasing current pulses (2s duration, starting at -50 pA with 5 pA increments). The Sag index was measured during 1000 ms current steps, with a peak voltage V_peak_ = -100 mV. Action potential (AP) and after-hyperpolarization (AHP) characteristics were measured during 1000 ms current steps near spike threshold. Mean max firing and spike adaptation properties were measured during 1000 ms current steps at the maximum depolarizing current that preceded the depolarization block. The range of current steps depended on input resistance and was set for each cell to cover voltage responses from -100 mV to the threshold to maximum firing. Spontaneous excitatory post-synaptic currents (sEPSCs) were evaluated in voltage-clamp at -50 mV.

### Immunohistochemistry

For tissue perfused with PFA, sections were washed 3 times for 10 minutes in phosphate-buffered saline (PBS) and blocked in a solution comprising PBS, 0.4% Triton X-100, and 5% normal goat or donkey serum (depending on host species) for 1 hr at room temperature. Sections were then incubated overnight (16-18 hours at 4 °C) in blocking solution with primary antibodies (see Supplementary Table 3). Next, sections were washed 3 times in PBS for 10 min and incubated with secondary antibodies (see Supplementary Table 3) in blocking solution for 1 hr at ambient temperature. Sections were washed again in PBS and incubated with Hoescht 33258 (pentahydrate (bis-benzimide); Dako, H1398) or 4’,6-diamidino-2-phenylindole (DAPI; Thermo Fisher Scientific, 62248) to visualize cell nuclei. Finally, sections were washed in PBS, mounted, and coverslipped on Superfrost Plus glass slides (Thermo Fisher Scientific) with Fluoromount G mounting medium (Thermo Fisher Scientific).

For tissue perfused with SSaCSF, sections were drop-fixed in 4% PFA in PBS overnight at 4 °C, then washed in PBS, permeabilized with 0.3% Triton X-100, and incubated with streptavidin (see Supplementary Table 3) overnight. Slices were then incubated in blocking solution at room temperature for at least 2 hours. Primary antibodies (see Supplementary Table 3) were diluted in carrier buffer containing 0.5% Triton X-100 for 3 hours at room temperature. After 3 PBS washes for 15 min each, the tissue was incubated with secondary antibodies (see Supplementary Table 3). After incubation, slices were incubated in DAPI (Millipore Sigma, D9542), washed multiple times, cryopreserved in 30% sucrose, and sectioned at 70 μm using a freezing microtome (Microm). Slices were then mounted on Superfrost Plus glass slides with Mowiol mounting medium.

### Microscopy

Images taken for quantitative cell counting were acquired using an Axio Imager Z1 epifluorescence microscope (Zeiss) equipped with an Apotome module for optical sectioning, or an Axioscan Z7 slide-scanning microscope (Zeiss). Images acquired with the Axio Imager Z1 used a Zeiss Axiocam HRm digital camera with Axiovision software (v.4.8.1.0) at a resolution of 0.2 x 0.2 µm per pixel. The objectives used were Zeiss EC Plan-Neofluar 5x/0.16 (420330-9901), EC Plan-Neofluar 10x/0.3 (420340-9901), and Plan Apochromat 20x/0.8 (420650-9901). Filter sets included Zeiss 49 DAPI (488049-9901-000), Zeiss HE eGFP (489038-9901-000), and Zeiss HQ Texas Red (000000-1114-462). Files were saved as Zeiss Vision Image (ZVI) and exported for analysis. Fluorescence images were acquired using the Axioscan Z7 with an AxioCam 712 mono (426560-9090-000) digital camera and ZEN 3.3 (blue edition) software at a resolution of 0.172 × 0.172 µm per pixel. The objective used was Zeiss Plan-Apochromat 20×/0.8 M27 (420650-9902-000). The Axioscan Z7 was equipped with a Viluma 7 LED light source and a Zeiss Filter set 112 HE LED w/module (489112-9110-400). Files were saved and exported as Carl Zeiss Image (CZI) files, then converted to Single Image Stack (SIS) files for analysis.

Images used to visualize cell population expression were acquired with the Axio Imager Z epifluorescence microscope using the above configuration, or a Mica microhub microscope (Leica Microsystems) in confocal mode using the LAS X software (version 6.2.3). The Mica microhub used a 20×/0.75 NA air objective (Leica HC PL APO CS2), and fluorescence emission was detected with Leica HyD FS (hybrid) detectors. Fluorophores were excited with 488 nm and 637 nm lasers using a T100 neutral density (ND) filter. Images were acquired with a pixel size of 75 × 75 nm, a pinhole size of 0.79 Airy units, and a frame acquisition time of 6.4 s. Sections containing biocytin-labeled cells were imaged using the Mica microhub in confocal mode. Z-stacks were acquired to span labelled cells using a 20×/0.75 NA air objective (Leica HC PL APO CS2) with a z-step size of 0.5 µm. The confocal pinhole was set to 1 Airy unit. Emitted fluorescence was detected using Leica HyD FS (hybrid) detectors. Fluorophores were excited with a 488 nm laser (10 mW maximum output), with laser transmission set to 4.4% using an acousto-optical tunable filter (AOTF) and a T10 neutral density (ND) filter. Images were acquired at 976 × 820 pixels with a pixel size of 120.3 × 120.3 nm and a pixel dwell time of 1.5 µs.

### Postprocessing and statistical analysis

#### Image processing and cell counting

To aid visualization, brightness and contrast adjustments were applied uniformly across individual microscopy images using Adobe Photoshop (version 25.12.4) or Fiji (version 1.53). PV/NeuN cell counts for PFC sections were performed in 2 sections per cortical hemisphere spanning the anteroposterior axis, and acquired in both cortical layer 3 and 5/6 in 6 anatomically defined areas (area 9, 46, 8aD, 8aV, 11, and 47L). In independent analyses, 2 methods were used for PV/NeuN cell counting. For images acquired with the Axio Z1 Imager microscope, cell counts were obtained from 3 randomly selected counting frames of 150,000 µm^2^ (approximately 440×340 µm), and supervised counting was performed using the *Cell Counter* plugin in Fiji (version 1.53) (*84*). For images acquired with the Axioscan Z7 slide-scanning microscope, counting frames were manually drawn across the entire radial extent of the layer of interest, and supervised counting was performed using Arivis Pro (version 4.2.2, Zeiss). Identical methods were used for cell counting in the primary motor cortex (M1; area 4ab), which was chosen as a negative control because it lacks direct connections to PM (*21*, *22*). For lesion quantification, NeuN counts in PM were obtained from 3 consecutive sections beginning at the interaural +3.0 mm level. As early-life lesions could not always be fully delineated visually due to longitudinal tissue recovery, cell counts for the EL and control cohorts were performed in 3 counting frames spaced across the mediolateral extent of PM. For adult lesions in which the core was identifiable, cell counts were obtained from counting frames of varying area within the lesion core. For consistency in reporting, all density values were normalized to counting frame area and represented as counts per 150,000 µm^2^.

#### Stereological estimation of pulvinar volume

To further confirm the presence of early lesions, PM volume estimates in the control and EL cohorts were obtained using the Cavalieri principle. In brief, calbindin-labeled sections, which accurately delineate PM boundaries, were sampled across the full anteroposterior extent of the pulvinar. PM volume was calculated by the sum of volumes for each section (PM area x 160 μm spacing), and corrected for linear shrinkage using a factor of 0.801 as previously defined in marmoset fixed sections (*85*).

#### Quantification of VGLUT2 puncta

The magnitude of VGLUT2 fluorescence surrounding PV+ cells was calculated by measuring fluorescence intensity in the perisomatic region. PV+ cells in layer 3 of 3 individual PFC areas (46, 47L, and 11) were examined at 40 x magnification (*n* = 13-22 cells per area). Analysis was performed using the wand tracing tool in Fiji, as previously described (*86*). In brief, the cell membrane was selected and its size was expanded by 0.5 μm, then the XOR option was used to restrict this mask to the edges of the perisoma. Perisomatic puncta were then quantified by subtracting the background and applying Gaussian blurring to the ROI using a sigma value of 1. Intensity values were measured across the perisomatic region and expressed as intensity per µm^2^.

#### Diffusivity analysis and surface mapping

Diffusivity maps were analyzed using voxel-based and ROI-based approaches. For the voxel-based approach, FA maps were first registered to a template space and thresholded to exclude voxels that putatively contained white matter (FA > 0.2) or cerebrospinal fluid (FA < 0.05). Next, pairwise statistical maps were produced for each comparison of age (6 mo., adult) and condition (control, EL) by applying two-sample voxel-wise t-tests with the *3dttest++* tool in AFNI (*87*). Thresholding was then applied to include only statistically significant voxels (*p* < 0.001), and clustering was performed for visualization purposes using a minimum of 200 contiguous, face-touching voxels. Finally, surface maps were generated by projecting the maximum mean difference values obtained from gray matter within clusters using AFNI *3dVol2Surf* and surface templates from the Marmoset Brain Mapping project (version 3.0) (*88*). For the ROI-based approach, the digital Paxinos atlas was registered to individual FA maps. Voxels were thresholded where FA < 0.05 or > 0.2, and the mean of all non-zero voxels was computed for cortical regions of interest. Cortical areas examined included areas 9, 46D, 46V, 8aV, 8aD, 8b, 11, 13a, 13b, 13M, 13L, 45, 47L, 47M, 47O, 25, 32V, 32, 10 and 4ab (M1).

#### Reversal learning analysis

Performance in the reversal learning task was primarily measured by errors to criterion, defined as the number of errors an animal committed during a task session. In addition, learning curve analysis was used to categorize trials within each session into discrete stages of learning. Using a previously described method (*38*, *39*, *89*, *90*), binary performance data from each reversal learning experiment were input into a recursive algorithm that detected ‘change points’ in the frequency of cumulative successes using χ^2^ tests. Trials were designated as change points if the frequency of cumulative successes at that point deviated from chance, as determined by a criterion of *p* < 0.02 (logit = 1.7). The change points were then used to segregate the learning curve into 3 stages: ‘perseverative’, ‘learning’, and ‘achieving’. Classification of trials into each stage was based on whether the mean performance between change points was significantly under, within, or above chance, as assessed using 2-tailed binomial tests. Next, we categorized ‘perseverative errors’ and ‘regressive errors’ as the number of errors committed within the perseverative and achieving stages, respectively. Finally, to assess how positive and negative feedback influenced acquisition of stimulus–reward contingencies, win–stay and lose–shift strategies were quantified during the learning phase of each reversal task. Win–stay probability was calculated as the proportion of correct responses relative to incorrect responses on trials immediately following a rewarded trial. Conversely, lose–shift probability was defined as the proportion of correct responses relative to incorrect responses on trials immediately following a non-rewarded trial. These values were averaged across all sessions per animal.

#### DMTS analysis

Performance in the DMTS was primarily measured by the average success rate per delay condition. To examine the effect of delay on working memory performance, the averaged performance data per delay length for each animal were fit with an exponential decay function:

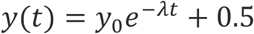

where *t* is time, *λ* is the decay constant and *y_0_* represents the zero-delay performance, or the estimated success rate at a delay of 0 s. The asymptote was constrained to the absolute chance level (i.e. 50%) to quantify ‘time to chance’, a measure of maximum maintenance capacity that we defined as the delay threshold where performance reached chance level according to a 1-tailed binomial test applied to the sample size of 0.5 – 4 s delays (*n* = 400 trials). The estimated zero-delay performance was used to quantify an animal’s baseline ability to successfully follow the task rule. ‘Half-life’ was calculated from the curve to describe the relationship between maintenance and delay and was defined as half the duration between maximal performance and chance. For visualization purposes, the decay function presented was the mean of the individual decay curves calculated for each animal, with the upper and lower boundaries representing standard deviation. Plots compare performance to the calculated chance threshold.

#### Spectral power analysis

For each recording session, power analysis was performed on clean, uninterrupted segments of the ECoG signal lasting at least 10 s. Using tools provided by MNE-Python (version 1.7) (*91*), power spectral densities for each segment were estimated with a multitaper method using 4 Slepian tapers, 624 ms windows, and a time-half-bandwidth product of 2.5. Session averages were calculated as the mean power spectral densities across all segments within a recording session. Absolute power values were then calculated for each frequency band of interest by estimating the area under the power spectral density curve using Simpson’s rule.

The frequency bands of interest were defined as follows: delta (0.5 – 4 Hz), theta (4 – 8 Hz), alpha (8 – 12 Hz), beta (12 – 30 Hz), and gamma (30 – 40 Hz). To normalize observations per individual session, relative power was then calculated as the percentage of absolute power per frequency band relative to the other bands. For visualization purposes, raw power spectra were log-transformed and expressed as decibels.

#### Patch clamp electrophysiology analysis

All electrophysiological analyses were conducted in Clampfit 10.7 (Molecular Devices) and processed in Excel (Microsoft) and MATLAB 2025a (MathWorks). RMP was calculated as the linearly extrapolated intersection point along the voltage ramp or as the mean voltage in I = 0 whole-cell recordings. Time constant τ was determined by fitting the mean voltage response to hyperpolarizing current injections with a single exponential function. R_in_ was determined by taking the slope of a linear regression of the change in voltage in response to increasing current pulses around the RMP, and the capacitance was given by C_in_ = τ/R_in_. The sag was determined by measuring the maximum hyperpolarization voltage deflection (V_peak_) from baseline (V_rest_) at the onset of the current pulse, and the steady-state voltage deflection (V_ss_) in the last 200 ms of the current pulse, with the sag index = (V_rest_ – V_ss_)/(V_rest_ – V_peak_). AP threshold was measured at the point at which the slope dV/dt exceeds 10 mV/ms, AP width (ms) at half the maximum voltage amplitude relative to threshold, and max up-/down-stroke as the max/min of the slope dV/dt, respectively. AHP amplitude and time were measured at the peak hyperpolarization voltage deflection relative to AP threshold. The maximum firing frequency was measured as the mean frequency during the maximum 1000 ms train preceding depolarization block, while the accommodation ratio was measured as the ratio of the final (last 3 spikes) to the initial (first 3 spikes) instantaneous frequency. sEPSCs were detected via template, with amplitude, rise, and decay computed from the mean sEPSC waveform of all detected events for each cell. A total of *n* = 51 control cells (*n* = 2 animals) and 48 EL cells (*n* = 2 animals) were recorded and analyzed. No data were excluded based on value, though some end-points could not be assessed for a given cell if the recording became unstable and the protocol was not reached.

#### Cell morphology reconstruction and quantification

Raw image stacks were imported into Zeiss Arivis Pro (version 4.2.2), and intensity was rescaled using a single optimal value for each image set by manually cropping the upper limit of the 8-bit histogram to ∼95–99.5 %. Somata were segmented using Blob Finder (diameter = 9–16 µm according to cell size, probability threshold = 65 %, split sensitivity = 90 %, normalization = ‘first time point’) followed by watershed Region Grower (threshold = 104) operations with manual curation. Somata volumes were exported directly from the object table. Primary neurites were counted manually in 2D and 3D views. The following exclusion conditions for individual cells were set prior to analysis: somata resides in underlying white matter; somata is within 15 µm of tissue damage or significant autofluorescent debris; or somata exhibits apoptotic or irregular morphology. 3 cells were excluded based on these criteria, and the remaining 68 cells (control: *n* = 37, lesion: *n* = 31) were used for subsequent analysis.

#### Statistical testing

The significance criterion for all tests besides MRI statistical mapping was set at *p* = 0.05. Due to constraints imposed by the small sample of nonhuman primates used in experiments, pairwise comparisons were conducted using permutation tests implemented in the Python package SciPy (version 1.17.1) (*92*). Monte Carlo permutations with 10,000 resamples were used to estimate p-values where the number of combinations exceeded 100,000.

Comparisons between multiple groups were conducted using a Kruskal-Wallis test with Dunn’s test for multiple comparisons, implemented in Prism 10 (GraphPad). For the DMTS analysis, trial-level performance was modeled using a generalized linear mixed model with a binomial distribution and logit link function using the *glmer* package in R, according to the following equation (in *glmer* notation):

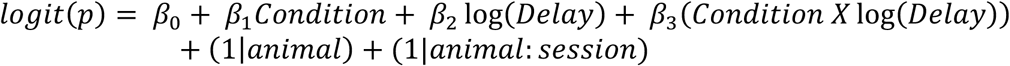

where *p* is the binary trial response (1 or 0), *Condition* is the animal grouping (i.e. control, EL, AD), *log(Delay)* is the linearized delay time, and both animal and session are treated as random intercepts to account for repeated observations. *Condition* was allocated as the reference group to visualize the effect of lesion conditions on response. The significance of fixed effects were evaluated with Wald *Z*-tests, and coefficients were displayed with 95% confidence intervals.

## Acknowledgements

The authors thank all members of the Bourne laboratory for helpful discussions; the McBain laboratory for technical assistance with patch clamp experiments and tissue processing; the Monash Animal Research Platform, the NIH Central Animal Facility, and Xianfeng (Lisa) Zhang for providing care and support for research animals; Monash Biomedical Imaging; the Neurophysiology Imaging Facility; the Monash Instrumentation Facility; the NIMH Section on Instrumentation; and the Systems Neuroscience Imaging Resource.

---

**Supplementary Figure 1.**
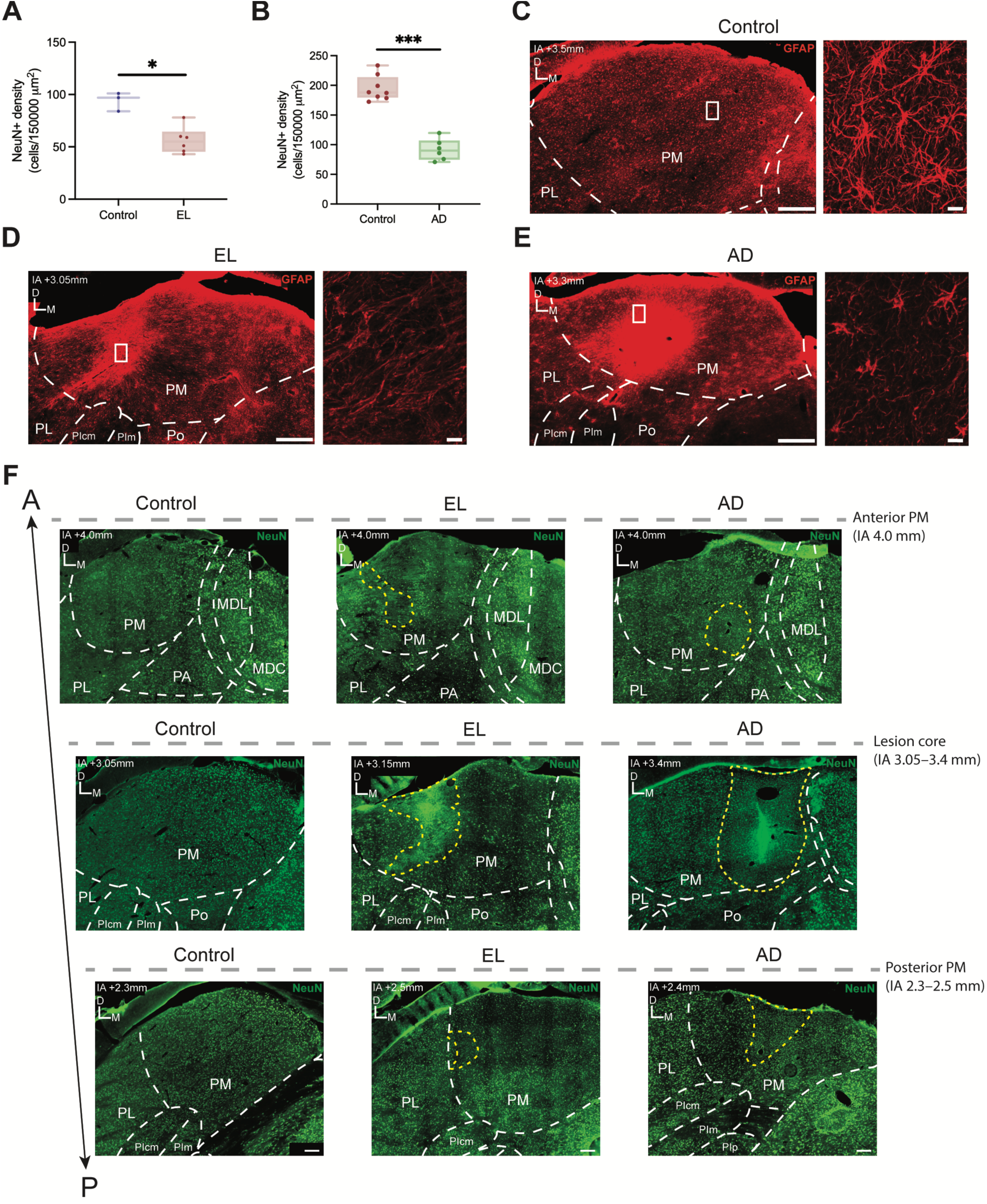
Extended anatomical assessment of PM lesions sustained in early life or adulthood. Reductions in NeuN+ cell density after **A.** EL and **B.** AD. **C-E.** Qualitative assessment of reactive astrogliosis within lesion cores, depicting formation of a glial scar within the lesion core and reactive astrocytes in the penumbral area. Scale bar = 500 µm, 10 µm. **F.** Representative sections along the anteroposterior axis of animals from control, EL, and AD conditions showing the predominant restriction of lesions to PM. Scale bar = 200 µm. PM: medial pulvinar; PL: lateral pulvinar; PIcm: inferior pulvinar (caudomedial division); PIm: inferior pulvinar (medial division); Po: posterior nuclei; MDL: mediodorsal nucleus (lateral division), MDC: mediodorsal nucleus (caudal division). **\*** *p* < 0.05, **\*\*\*** *p* < 0.001.

**Supplementary Figure 2.**
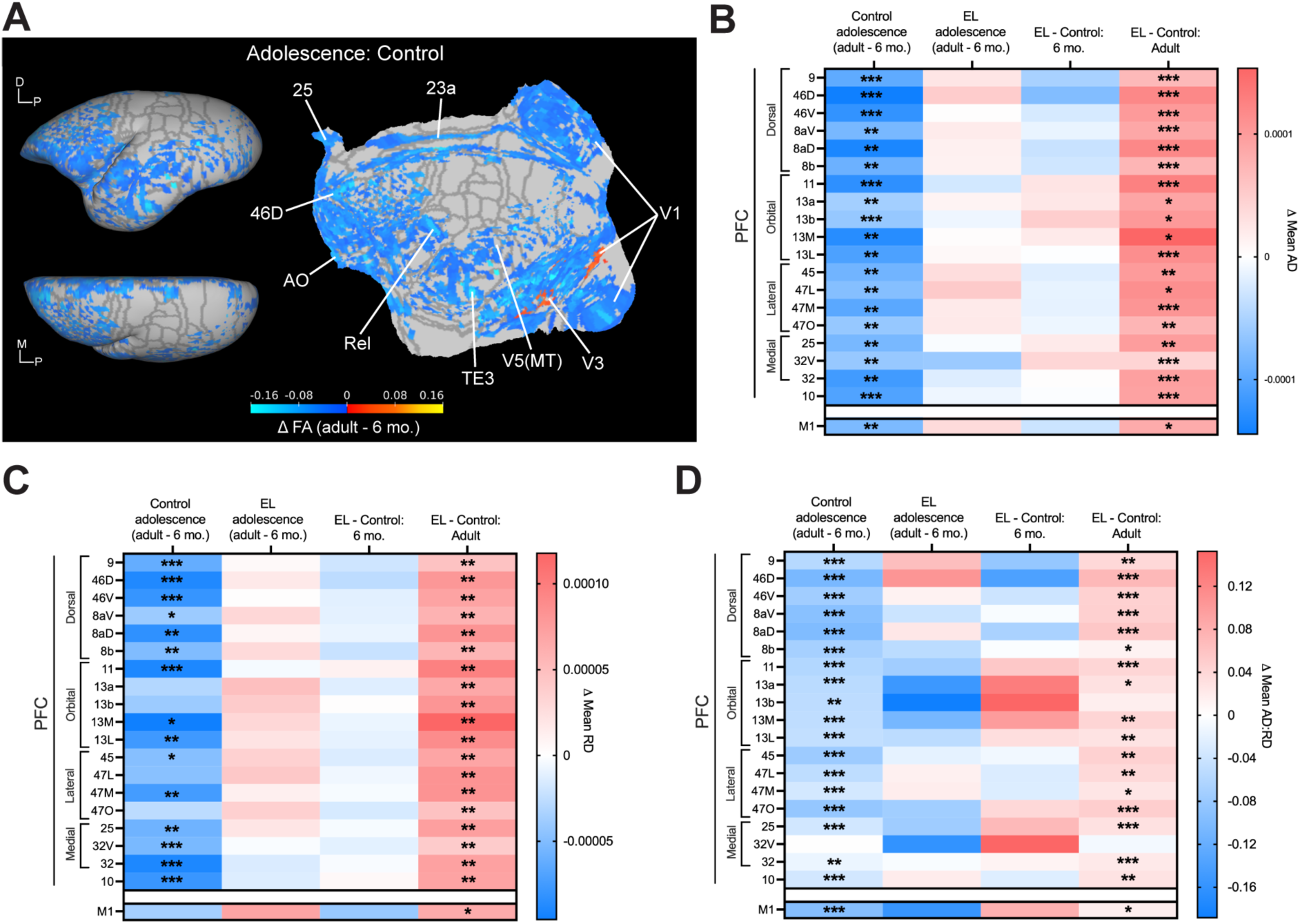
Extended data revealing changes in cortical diffusivity over adolescence in EL compared to control animals. **A.** Statistical flatmap and volume renderings of voxel-wise FA differences in cortical gray matter between adulthood (>18 mo.) and 6 mo. in controls, indicative of typical adolescent maturation. Lower values indicate lesser FA in adulthood relative to 6 mo. **B-D.** ROI analysis of mean axial diffusivity (AD), radial diffusivity (RD), and AD:RD ratio across constituent areas of the PFC and M1 across age and condition. **\*** *p* < 0.05, **\*\*** *p* < 0.01, **\*\*\*** *p* < 0.001.

**Supplementary Figure 3.**
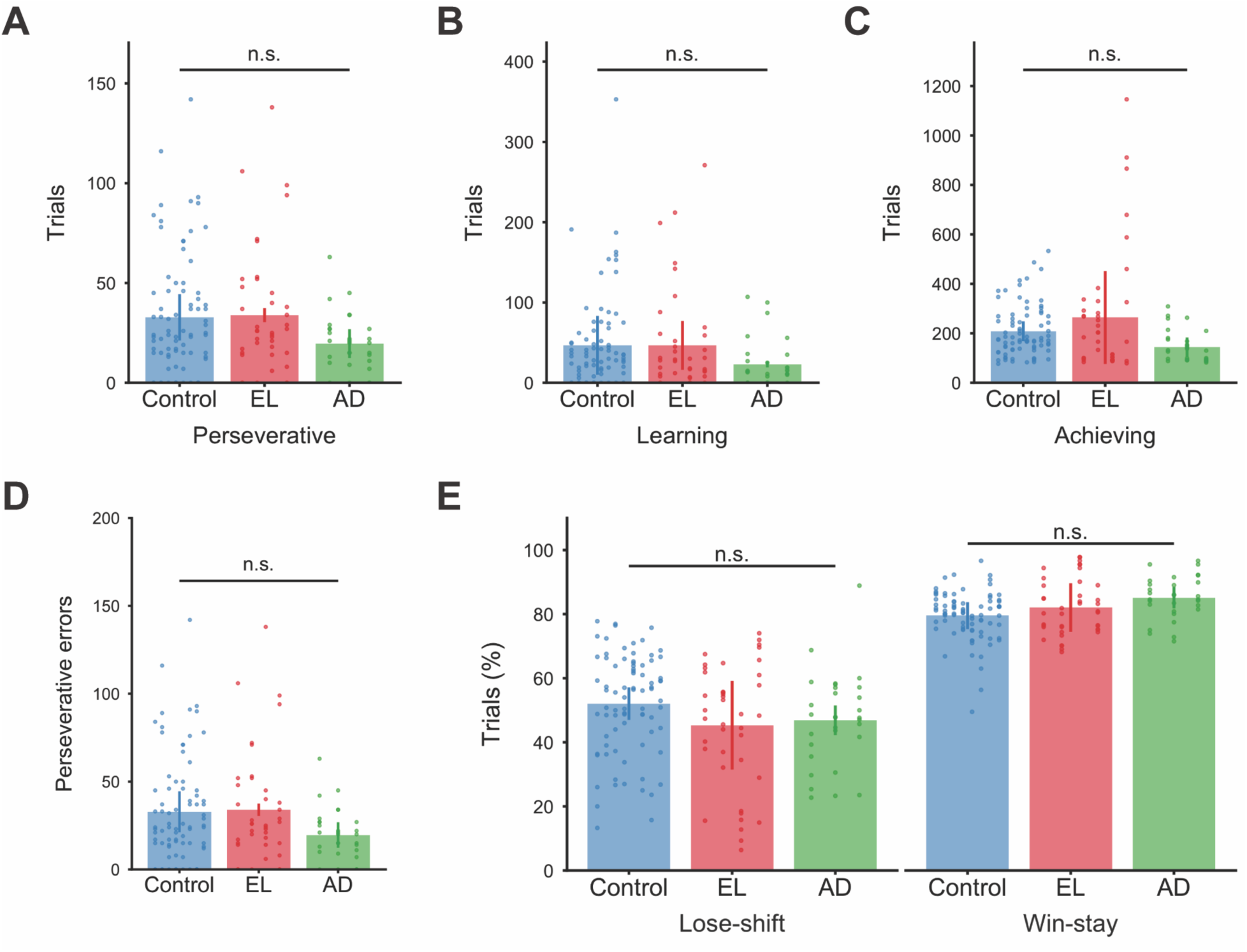
Extended characterization of cognitive flexibility during reversal learning following early life or adult PM lesions. **A-C.** Number of trials performed across control, EL, and AD conditions in the perseverative, learning, and achieving stages of reversal learning, as defined by learning curve analysis. **D.** Number of errors committed during the perseverative stage across conditions. **E.** The proportions of lose-shift and win-stay response strategies performed in reversal learning sessions across conditions. Percentages are calculated based on the possible trial outcomes given the previous trial. Data points in the overlaid vertical scatter plot lines represent individual session performance for each animal.

**Supplementary Figure 4.**
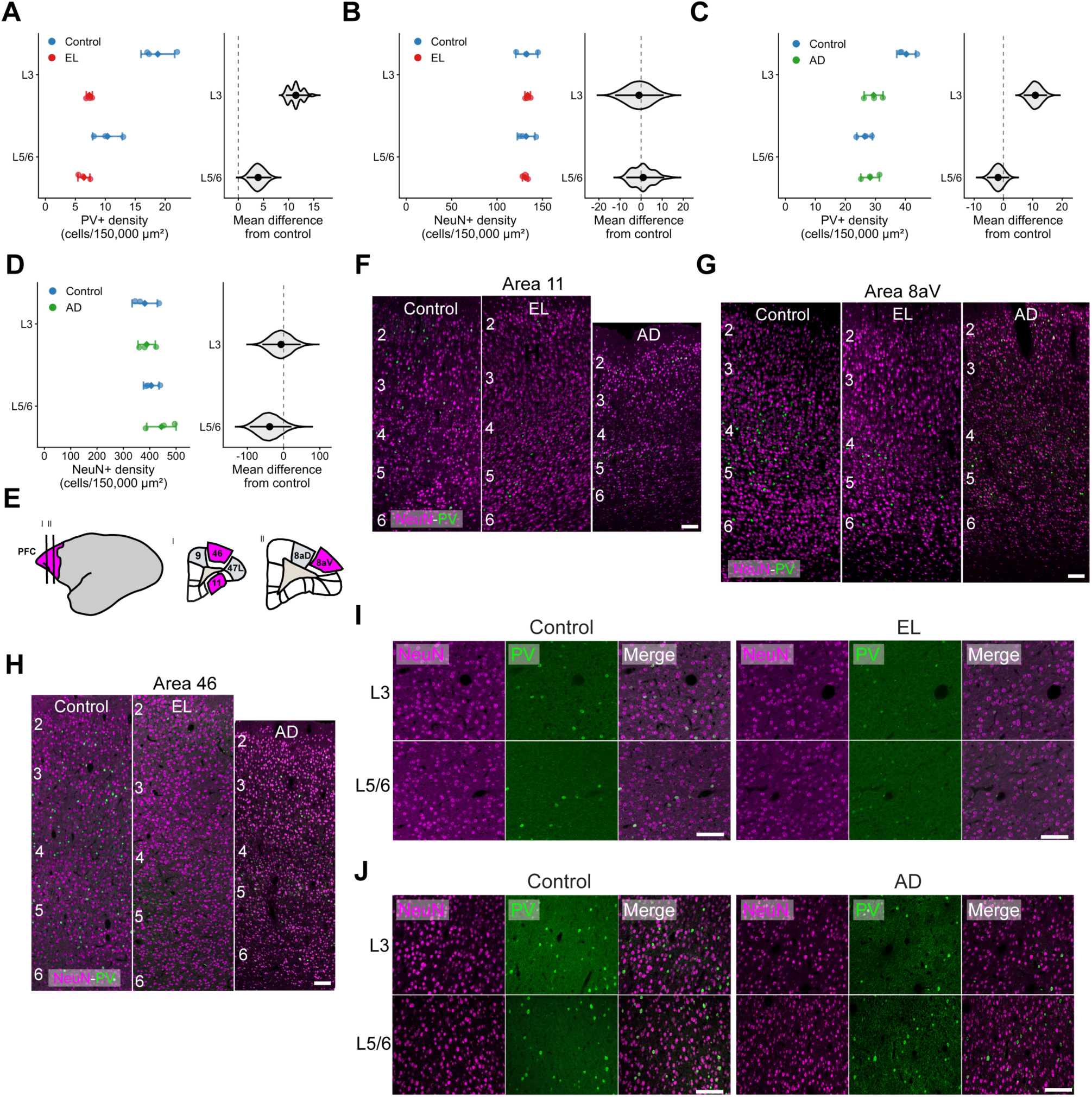
Extended assessment of PV and NeuN expression in PFC following early life or adult PM lesions. **A-D.** Gardner-Altman estimation plots of raw cell density counts from both PV and NeuN immunolabelling in control, EL, and AD tissue. As the microscopy methods differed between EL and AD tissue, each cohort was compared with an independent cohort of method-matched control tissue. Left plots display mean +/- SD, right plots display mean difference +/- 95% bootstrapped CI and the bootstrapped distribution of the mean difference. **E.** Schematic of the PFC sections used for cell counting in the current figure. **F-H.** Laminar expression of NeuN/PV immunolabelling in control, EL, and AD tissue across 3 representative PFC areas. Scale bar = 100 µm. **I, J.** Representative images used for cell counting of PV/NeuN immunolabelling in layers 3 and 5/6. Scale bar = 100 µm.

**Supplementary Figure 5.**
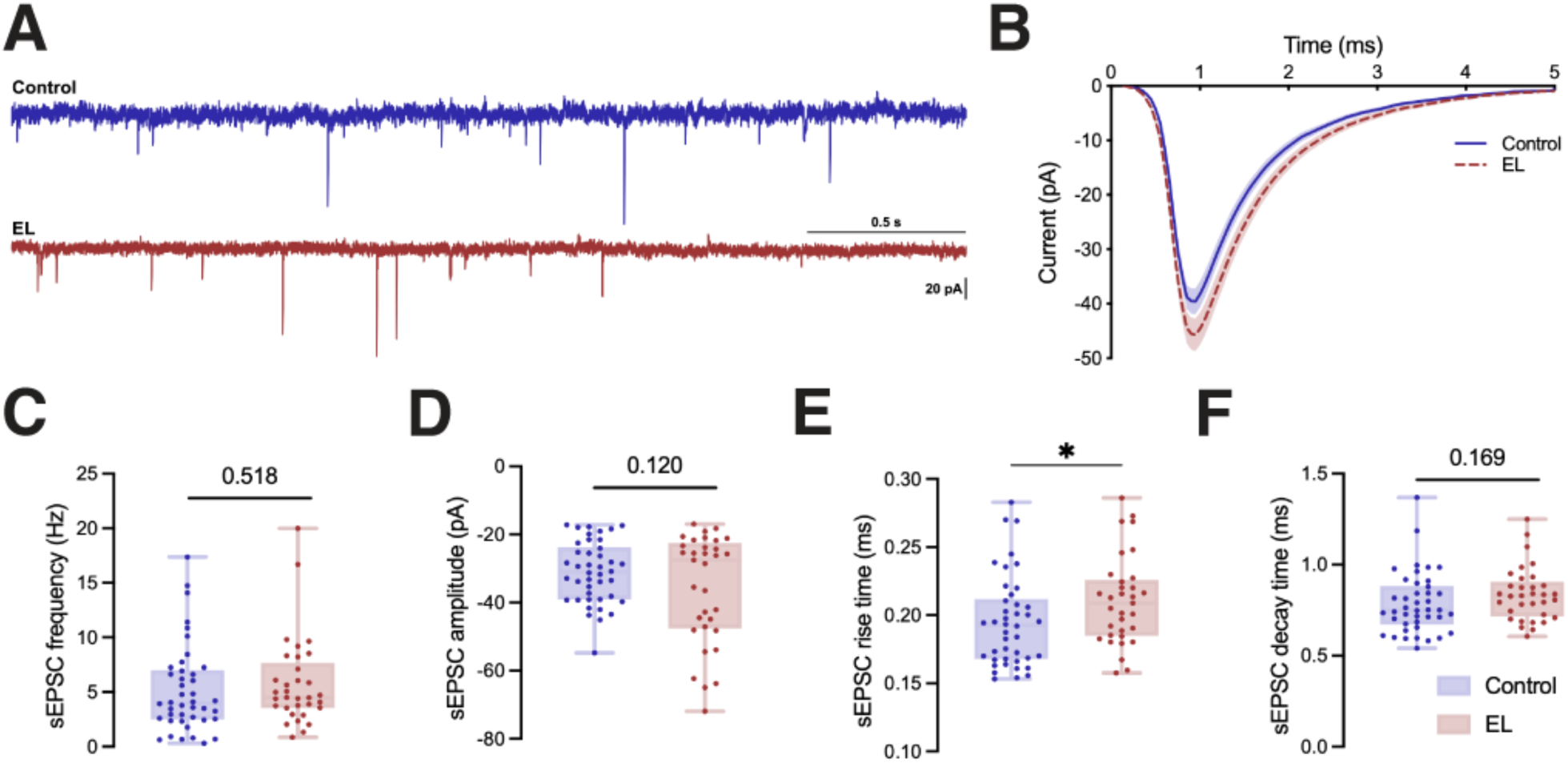
Preservation of sEPSCs in PV-class interneurons of the PFC following early PM lesions. **A.** Representative sEPSC trace comparing control and EL conditions. **B.** Temporal dynamics of recorded sEPSCs. Data displays the mean of individual cell responses +/- SEM. **C-F.** Electrophysiological characteristics of recorded sEPSCs comparing control and EL conditions. **\*** *p <* 0.05.

**Supplementary Table 1.**
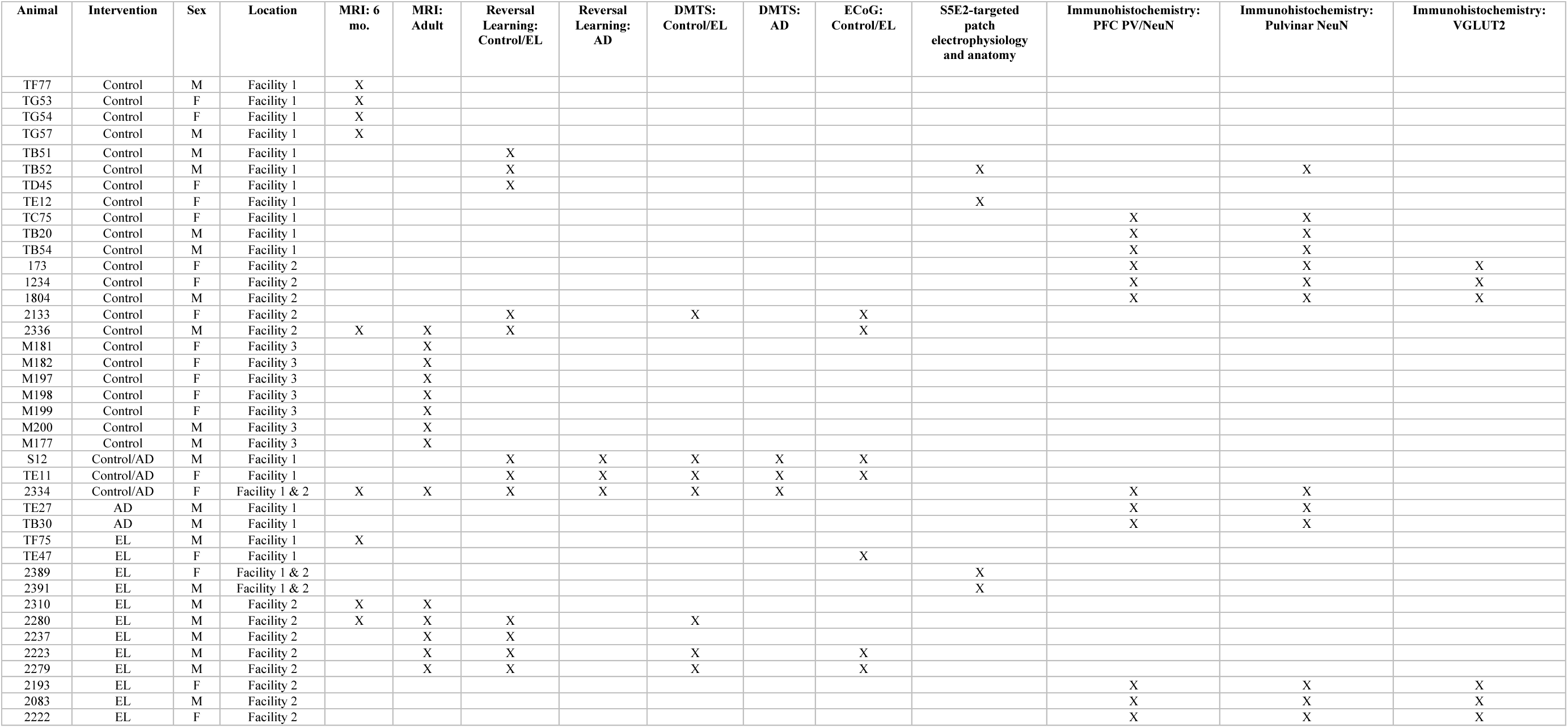
Experimental allocation of animals used in the study.

**Supplementary Table 2.** Electrophysiological parameters of S5E2+ cells.

| Parameter | Control | EL | Permutation test p-value |
| --- | --- | --- | --- |
| RMP (mV) | -62.0 ± 1.1 (n = 51) | -59.5 ± 1.0 (n = 47) | 0.099 |
| Spont. AP Frequency (Hz) | 1.46 ± 0.64 (n = 44) | 5.45 ± 1.94 (n = 37) | 0.038 (*) |
| Tau (ms) | 10.6 ± 0.5 (n = 52) | 13.6 ± 1.0 (n = 47) | 0.001 (**) |
| R<sub>in</sub> (MΩ) | 124.6 ± 6.5 (n = 50) | 187.2 ± 13.6 (n = 45) | 0.0005 (****) |
| C<sub>in</sub> (pF) | 95.2 ± 6.7 (n = 50) | 80.2 ± 5.7 (n = 45) | 0.087 |
| Sag (mV) | 7.40 ± 0.45 (n = 44) | 7.08 ± 0.48 (n = 48) | 0.633 |
| Sag Index | 0.77 ± 0.01 (n = 51) | 0.78 ± 0.01 (n = 48) | 0.568 |
| Max. Frequency (Hz) | 249 ± 9 (n = 51) | 216 ± 11 (n = 47) | 0.022 (*) |
| Inst. Frequency (Hz) | 313 ± 10 (n = 51) | 286 ± 13 (n = 47) | 0.113 |
| Accommodation | 0.76 ± 0.01 (n = 51) | 0.72 ± 0.02 (n = 47) | 0.043 (*) |
| AP Threshold (mV) | -41.5 ± 0.7 (n = 51) | -40.7 ± 0.7 (n = 47) | 0.452 |
| AP Amplitude (mV) | 47.8 ± 1.1 (n = 51) | 49.4 ± 1.4 (n = 47) | 0.360 |
| AP Width (ms) | 0.36 ± 0.01 (n = 51) | 0.43 ± 0.02 (n = 47) | 0.002 (**) |
| Max Upstroke (mV/ms) | 253.6 ± 8.8 (n = 51) | 234.8 ± 8.6 (n = 47) | 0.128 |
| Max Downstroke (mV/ms) | -177.4 ± 7.8 (n = 51) | -154.5 ± 8.2 (n = 47) | 0.045 (*) |
| AHP Amplitude (mV) | -19.9 ± 0.7 (n = 48) | -20.1 ± 0.6 (n = 47) | 0.883 |
| AHP Time (ms) | 1.93 ± 0.14 (n = 48) | 3.29 ± 0.98 (n = 47) | 0.039 (*) |
| sEPSC Frequency (Hz) | 5.17 ± 0.62 (n = 42) | 5.78 ± 0.70 (n = 33) | 0.519 |
| sEPSC Amplitude (pA) | -31.1 ± 1.4 (n = 42) | -35.7 ± 2.8 (n = 33) | 0.120 |
| sEPSC Rise Time (ms) | 0.195 ± 0.005 (n = 42) | 0.211 ± 0.006 (n = 33) | 0.047 (*) |
| sEPSC Decay Time (ms) | 0.785 ± 0.026 (n = 42) | 0.837 ± 0.026 (n = 33) | 0.169 |

**Supplementary Table 3.** Primary and secondary antibodies used for immunohistochemistry.

| Antigen | Figure abbreviation | Host species | Dilution | Source | Catalog ID | RRID |
| --- | --- | --- | --- | --- | --- | --- |
| **Primary** |  |  |  |  |  |  |
| Neuronal nuclear antigen | NeuN | Rabbit | 1:1000 | Millipore-Sigma | ABN78 | AB_10807945 |
| Parvalbumin | PV | Rabbit | 1:1000 | Swant | PV28 | AB_2315235 |
| Parvalbumin | PV | Mouse | 1:1000 | Swant | PV235 | AB_3698492 |
| Parvalbumin | PV | Mouse | 1:1000 | Millipore-Sigma | P3088 | AB_477329 |
| Glial fibrillary acidic protein | GFAP | Mouse | 1:1000 | Millipore-Sigma | MAB360 | AB_11212597 |
| Vesicular glutamate transporter 2 | VGLUT2 | Mouse | 1:500 | Millipore-Sigma | MAB5504 | AB_2187552 |
| Calbindin | Calb | Mouse | 1:1000 | Swant | CB300 | AB_10000347 |
| DsRed | S5E2 | Rabbit | 1:1000 | Takara | 632496 | AB_10013483 |
| **Secondary** |  |  |  |  |  |  |
| Donkey anti-mouse Alexa Fluor 488 | - | Donkey | 1:800 | Thermo Fisher | A21202 | AB_141607 |
| Donkey anti-mouse Alexa Fluor 647 | - | Donkey | 1:800 | Thermo Fisher | A31571 | AB_162542 |
| Donkey anti-rabbit Alexa Fluor 488 | - | Donkey | 1:800 | Thermo Fisher | A21206 | AB_2535792 |
| Donkey anti-rabbit Alexa Fluor 594 | - | Donkey | 1:800 | Thermo Fisher | A21207 | AB_141637 |
| Goat anti-rabbit Alexa Fluor 488 | - | Goat | 1:800 | Thermo Fisher | A11008 | AB_143165 |
| Goat anti-rabbit Alexa Fluor 488 | - | Goat | 1:1000 | Thermo Fisher | A11034 | AB_2576217 |
| Goat anti-mouse Alexa Fluor 647 | - | Goat | 1:800 | Thermo Fisher | A21235 | AB_2535804 |
| Goat anti-rabbit Alexa Fluor 555 | - | Goat | 1:1000 | Thermo Fisher | A21429 | AB_2535850 |
| **Other conjugate** |  |  |  |  |  |  |
| Streptavidin<br>Alexa Fluor<br>555 | - | - | 1:500 | Thermo<br>Fisher | S32355 | AB_2571525 |

## References

1. J. Smucny, S. J. Dienel, D. A. Lewis, C. S. Carter, Mechanisms underlying dorsolateral prefrontal cortex contributions to cognitive dysfunction in schizophrenia. Neuropsychopharmacol. 47, 292–308 (2022).

2. A. F. T. Arnsten, Prefrontal Cortical Network Connections: Key Site of Vulnerability in Stress and Schizophrenia. Int J Dev Neurosci 29, 215–223 (2011).

3. N. Gogtay, J. N. Giedd, L. Lusk, K. M. Hayashi, D. Greenstein, A. C. Vaituzis, T. F. Nugent, D. H. Herman, L. S. Clasen, A. W. Toga, J. L. Rapoport, P. M. Thompson, Dynamic mapping of human cortical development during childhood through early adulthood. Proc Natl Acad Sci U S A 101, 8174–8179 (2004).

4. Z. Petanjek, M. Judaš, G. Šimić, M. R. Rašin, H. B. M. Uylings, P. Rakic, I. Kostović, Extraordinary neoteny of synaptic spines in the human prefrontal cortex. Proc Natl Acad Sci U S A 108, 13281–13286 (2011).

5. K. J. Burman, L. L. Lui, M. G. P. Rosa, J. A. Bourne, Development of non-phosphorylated neurofilament protein expression in neurones of the New World monkey dorsolateral frontal cortex. European Journal of Neuroscience 25, 1767–1779 (2007).

6. N. S. Hosseini Fin, A. Yip, J. T. Scott, L. Teo, J. Homman-Ludiye, J. A. Bourne, Developmental dynamics of marmoset prefrontal cortical SST and PV interneuron networks highlight primate-specific features. Development 152, dev204254 (2025).

7. L. A. Glantz, D. A. Lewis, Decreased Dendritic Spine Density on Prefrontal Cortical Pyramidal Neurons in Schizophrenia. Arch Gen Psychiatry 57, 65–73 (2000).

8. A. Sekar, A. R. Bialas, H. de Rivera, A. Davis, T. R. Hammond, N. Kamitaki, K. Tooley, J. Presumey, M. Baum, V. Van Doren, G. Genovese, S. A. Rose, R. E. Handsaker, Schizophrenia Working Group of the Psychiatric Genomics Consortium, M. J. Daly, M. C. Carroll, B. Stevens, S. A. McCarroll, Schizophrenia risk from complex variation of complement component 4. Nature 530, 177–183 (2016).

9. D. W. Volk, J. R. Edelson, D. A. Lewis, Altered expression of developmental regulators of parvalbumin and somatostatin neurons in the prefrontal cortex in schizophrenia. Schizophr Res 177, 3–9 (2016).

10. O. Marín, Parvalbumin interneuron deficits in schizophrenia. Eur Neuropsychopharmacol 82, 44–52 (2024).

11. J. F. Enwright, S. Sanapala, A. Foglio, R. Berry, K. N. Fish, D. A. Lewis, Reduced Labeling of Parvalbumin Neurons and Perineuronal Nets in the Dorsolateral Prefrontal Cortex of Subjects with Schizophrenia. Neuropsychopharmacology 41, 2206–2214 (2016).

12. T. Spadory, A. Duque, L. D. Selemon, Spatial-temporal topography in neurogenesis of the macaque thalamus. Brain Struct Funct 227, 1673–1682 (2022).

13. M. Alhesain, A. Alzu’bi, N. Sankar, C. Smith, J. Kerwin, R. Laws, S. Lindsay, G. J. Clowry, Development of the early fetal human thalamus: from a protomap to emergent thalamic nuclei. Front Neuroanat 19, 1530236 (2025).

14. V. J. Sydnor, J. Bagautdinova, B. Larsen, M. J. Arcaro, D. M. Barch, D. S. Bassett, A. F. Alexander-Bloch, P. A. Cook, S. Covitz, A. R. Franco, R. E. Gur, R. C. Gur, A. P. Mackey, K. Mehta, S. L. Meisler, M. P. Milham, T. M. Moore, E. J. Müller, D. R. Roalf, T. Salo, G. Schubiner, J. Seidlitz, R. T. Shinohara, J. M. Shine, F.-C. Yeh, M. Cieslak, T. D. Satterthwaite, Human thalamocortical structural connectivity develops in line with a hierarchical axis of cortical plasticity. Nat Neurosci 28, 1772–1786 (2025).

15. T. Guillamón-Vivancos, M. Aníbal-Martínez, L. Puche-Aroca, F. J. Martini, G. López-Bendito, Sensory modality-specific wiring of thalamocortical circuits. Nat. Rev. Neurosci. 26, 623–641 (2025).

16. T. E. Faust, G. Gunner, D. P. Schafer, Mechanisms governing activity-dependent synaptic pruning in the developing mammalian CNS. Nat Rev Neurosci 22, 657–673 (2021).

17. I.-C. Mundinano, D. M. Fox, W. C. Kwan, D. Vidaurre, L. Teo, J. Homman-Ludiye, M. A. Goodale, D. A. Leopold, J. A. Bourne, Transient visual pathway critical for normal development of primate grasping behavior. Proceedings of the National Academy of Sciences 115, 1364–1369 (2018).

18. L. J. Benoit, E. S. Holt, L. Posani, S. Fusi, A. Z. Harris, S. Canetta, C. Kellendonk, Adolescent thalamic inhibition leads to long-lasting impairments in prefrontal cortex function. Nat Neurosci 25, 714–725 (2022).

19. S.-S. Yang, Q. He, X. Gu, S. Liu, W. Ke, L. Chen, B. Li, Y. Shu, W.-J. Gao, Transient Inhibition of the Mediodorsal Thalamus During Early Adolescence Induces Hypofrontality and Social Memory Deficits in Young Adulthood. Biological Psychiatry Global Open Science 5, 100486 (2025).

20. D. Petersen, R. Raudales, A. K. Silva, C. Kellendonk, S. Canetta, Adolescent Thalamoprefrontal Inhibition Leads to Changes in Intrinsic Prefrontal Network Connectivity. eNeuro 11, ENEURO.0284–24.2024 (2024).

21. M. A. Córdoba-Claros, P. Rubio-Garrido, R. R. M. de Lima, P. L. A. G. Morais, E. S. do Nascimento, J. S. Cavalcante, F. Clascá, Projection Motifs and Wiring Logic of Medial Pulvinar Thalamocortical Axons in the Marmoset Monkey. J. Neurosci. 45 (2025).

22. J. Homman-Ludiye, I. C. Mundinano, W. C. Kwan, J. A. Bourne, Extensive Connectivity Between the Medial Pulvinar and the Cortex Revealed in the Marmoset Monkey. Cereb Cortex 30, 1797–1812 (2020).

23. K.-A. Dorph-Petersen, D. A. Lewis, Postmortem structural studies of the thalamus in schizophrenia. Schizophr Res 180, 28–35 (2017).

24. W. Byne, J. Fernandes, V. Haroutunian, D. Huacon, S. Kidkardnee, J. Kim, A. Tatusov, U. Thakur, G. Yiannoulos, Reduction of right medial pulvinar volume and neuron number in schizophrenia. Schizophrenia Research 90, 71–75 (2007).

25. A. S. Huang, B. P. Rogers, J. M. Sheffield, M. E. Jalbrzikowski, A. Anticevic, J. U. Blackford, S. Heckers, N. D. Woodward, Thalamic Nuclei Volumes in Psychotic Disorders and in Youths With Psychosis Spectrum Symptoms. Am J Psychiatry 177, 1159–1167 (2020).

26. D. Cobia, C. Rich, M. J. Smith, P. Engel Gonzalez, W. Cronenwett, J. G. Csernansky, L. Wang, Thalamic Shape Abnormalities Differentially Relate to Cognitive Performance in Early-Onset and Adult-Onset Schizophrenia. Front Psychiatry 13, 803234 (2022).

27. J. Penner, E. A. Osuch, B. Schaefer, J. Théberge, R. W. J. Neufeld, R. S. Menon, N. Rajakumar, J. A. Bourne, P. C. Williamson, Higher order thalamic nuclei resting network connectivity in early schizophrenia and major depressive disorder. Psychiatry Res Neuroimaging 272, 7–16 (2018).

28. A. Alzu’bi, J. Homman-Ludiye, J. A. Bourne, G. J. Clowry, Thalamocortical Afferents Innervate the Cortical Subplate much Earlier in Development in Primate than in Rodent. Cereb Cortex 29, 1706–1718 (2019).

29. J. Homman-Ludiye, W. C. Kwan, M. J. de Souza, J. A. Bourne, Full: Ontogenesis and development of the nonhuman primate pulvinar. Journal of Comparative Neurology 526, 2870–2883 (2018).

30. S. J. Sawiak, Y. Shiba, L. Oikonomidis, C. P. Windle, A. M. Santangelo, H. Grydeland, G. Cockcroft, E. T. Bullmore, A. C. Roberts, Trajectories and Milestones of Cortical and Subcortical Development of the Marmoset Brain From Infancy to Adulthood. Cereb Cortex 28, 4440–4453 (2018).

31. C. Reveley, F. Q. Ye, R. B. Mars, D. Matrov, Y. Chudasama, D. A. Leopold, Diffusion MRI anisotropy in the cerebral cortex is determined by unmyelinated tissue features. Nat Commun 13, 6702 (2022).

32. J. A. Acosta-Franco, G. Little, C. Beaulieu, High resolution diffusion tensor imaging of the human cortex reveals non-linear trajectories over the healthy lifespan. Imaging Neuroscience 3, IMAG.a.115 (2025).

33. K. M. Lynch, R. P. Cabeen, A. W. Toga, Spatiotemporal patterns of cortical microstructural maturation in children and adolescents with diffusion MRI. Human Brain Mapping 45, e26528 (2024).

34. C. D. Kroenke, Using diffusion anisotropy to study cerebral cortical gray matter development. J Magn Reson 292, 106–116 (2018).

35. M. Ouyang, T. Jeon, A. Sotiras, Q. Peng, V. Mishra, C. Halovanic, M. Chen, L. Chalak, N. Rollins, T. P. L. Roberts, C. Davatzikos, H. Huang, Differential cortical microstructural maturation in the preterm human brain with diffusion kurtosis and tensor imaging. Proceedings of the National Academy of Sciences 116, 4681–4688 (2019).

36. R. Dias, T. W. Robbins, A. C. Roberts, Dissociable Forms of Inhibitory Control within Prefrontal Cortex with an Analog of the Wisconsin Card Sort Test: Restriction to Novel Situations and Independence from “On-Line” Processing. J Neurosci 17, 9285–9297 (1997).

37. R. Dias, T. W. Robbins, A. C. Roberts, Dissociation in prefrontal cortex of affective and attentional shifts. Nature 380, 69–72 (1996).

38. J. T. Scott, B. L. M. Vasquez, B. J. Stewart, D. D. Panacheril, D. K. J. Rajit, A. Y. Fan, J. A. Bourne, CalliCog is an open-source cognitive neuroscience toolkit for freely behaving nonhuman primates. Cell Reports Methods 5 (2025).

39. C. R. Gallistel, S. Fairhurst, P. Balsam, The learning curve: Implications of a quantitative analysis. Proceedings of the National Academy of Sciences 101, 13124–13131 (2004).

40. C. Basar-Eroglu, A. Brand, H. Hildebrandt, K. Karolina Kedzior, B. Mathes, C. Schmiedt, Working memory related gamma oscillations in schizophrenia patients. Int J Psychophysiol 64, 39–45 (2007).

41. C. Haenschel, R. A. Bittner, J. Waltz, F. Haertling, M. Wibral, W. Singer, D. E. J. Linden, E. Rodriguez, Cortical oscillatory activity is critical for working memory as revealed by deficits in early-onset schizophrenia. J Neurosci 29, 9481–9489 (2009).

42. G. Buzsáki, X.-J. Wang, Mechanisms of gamma oscillations. Annu Rev Neurosci 35, 203–225 (2012).

43. K. K. A. Cho, R. Hoch, A. T. Lee, T. Patel, J. L. R. Rubenstein, V. S. Sohal, Gamma rhythms link prefrontal interneuron dysfunction with cognitive inflexibility in Dlx5/6(+/-) mice. Neuron 85, 1332–1343 (2015).

44. M. Lagler, A. T. Ozdemir, S. Lagoun, H. Malagon-Vina, Z. Borhegyi, R. Hauer, A. Jelem, T. Klausberger, Divisions of Identified Parvalbumin-Expressing Basket Cells during Working Memory-Guided Decision Making. Neuron 91, 1390–1401 (2016).

45. A. Mukherjee, N. H. Lam, R. D. Wimmer, M. M. Halassa, Thalamic circuits for independent control of prefrontal signal and noise. Nature 600, 100–104 (2021).

46. K. Delevich, J. Tucciarone, Z. J. Huang, B. Li, The Mediodorsal Thalamus Drives Feedforward Inhibition in the Anterior Cingulate Cortex via Parvalbumin Interneurons. J Neurosci 35, 5743–5753 (2015).

47. K. N. Fish, G. D. Hoftman, W. Sheikh, M. Kitchens, D. A. Lewis, Parvalbumin-Containing Chandelier and Basket Cell Boutons Have Distinctive Modes of Maturation in Monkey Prefrontal Cortex. J Neurosci 33, 8352–8358 (2013).

48. S. L. Erickson, D. A. Lewis, Postnatal development of parvalbumin- and GABA transporter-immunoreactive axon terminals in monkey prefrontal cortex. Journal of Comparative Neurology 448, 186–202 (2002).

49. M. Moissidis, L. Abbasova, M. Selten, R. Alis, C. Bernard, Y. Domínguez-Canterla, F. Oozeer, S. Qin, A. Kelly, L. Mòdol, N. A. Vasistha, B. Jones, P. Dhami, K. Khodosevich, F. Hamid, P. Lavender, N. Flames, O. Marín, A postnatal molecular switch drives activity-dependent maturation of parvalbumin interneurons. Cell 188, 5555–5575.e26 (2025).

50. S. E. Canetta, E. S. Holt, L. J. Benoit, E. Teboul, G. M. Sahyoun, R. T. Ogden, A. Z. Harris, C. Kellendonk, Mature parvalbumin interneuron function in prefrontal cortex requires activity during a postnatal sensitive period. eLife 11, e80324 (2022).

51. J. Homman-Ludiye, J. A. Bourne, The medial pulvinar: function, origin and association with neurodevelopmental disorders. Journal of Anatomy 235, 507–520 (2019).

52. D. Vormstein-Schneider, J. D. Lin, K. A. Pelkey, R. Chittajallu, B. Guo, M. A. Arias-Garcia, K. Allaway, S. Sakopoulos, G. Schneider, O. Stevenson, J. Vergara, J. Sharma, Q. Zhang, T. P. Franken, J. Smith, L. A. Ibrahim, K. J. M astro, E. Sabri, S. Huang, E. Favuzzi, T. Burbridge, Q. Xu, L. Guo, I. Vogel, V. Sanchez, G. A. Saldi, B. L. Gorissen, X. Yuan, K. A. Zaghloul, O. Devinsky, B. L. Sabatini, R. Batista-Brito, J. Reynolds, G. Feng, Z. Fu, C. J. McBain, G. Fishell, J. Dimidschstein, Viral manipulation of functionally distinct interneurons in mice, non-human primates and humans. Nat Neurosci 23, 1629–1636 (2020).

53. N. D. Woodward, H. Karbasforoushan, S. Heckers, Thalamocortical dysconnectivity in schizophrenia. Am J Psychiatry 169, 10.1176/appi.ajp.2012.12010056 (2012).

54. M. Giraldo-Chica, B. P. Rogers, S. M. Damon, B. A. Landman, N. D. Woodward, Prefrontal-Thalamic Anatomical Connectivity and Executive Cognitive Function in Schizophrenia. Biological Psychiatry 83, 509–517 (2018).

55. L. D. Selemon, N. Zecevic, Schizophrenia: a tale of two critical periods for prefrontal cortical development. Transl Psychiatry 5, e623 (2015).

56. A. Schmitt, P. Falkai, S. Papiol, Neurodevelopmental disturbances in schizophrenia: evidence from genetic and environmental factors. J Neural Transm 130, 195–205 (2023).

57. J. Homman-Ludiye, J. A. Bourne, “The medial pulvinar: A crucial player in the development of the primate brain” in Factors Affecting Neurodevelopment (Academic Press, 2021; https://www.sciencedirect.com/science/chapter/edited-volume/pii/B9780128179864000304), pp. 347–357.

58. A. Lella, L. A. Antonucci, R. Passiatore, L. Bellantuono, P. Selvaggi, T. Popolizio, G. Di Sciascio, A. Saponaro, P. Ricci, M. Altamura, G. Blasi, A. Rampino, C. Vriend, V. D. Calhoun, K. Rootes-Murdy, A. L. Goldman, I. Baeza, J. Castro-Fornieles, G. Sugranyes, E. De la Serna, E. Pomarol-Clotet, M. Fatjó-Vilas, R. Salvador, A. Karuk, P. Fuentes-Claramonte, D. C. Glahn, A. L. Rodrigue, J. Blangero, L. Wang, T. Lee, K. E. Einenkel, S. Hamers, O. Gruber, A. Preda, Y.-C. Chung, S. Odkhuu, C. Vallée, P. Dazzan, M. Marcelis, S. Michielse, K. Brosch, F. Stein, I. Nenadić, B. Straube, F. Thomas-Odenthal, T. Kircher, S. Carruthers, S. L. Rossell, P. J. Sumner, T. E. Van Rheenen, C. Demro, I. S. Ramsay, S. R. Sponheim, R. Lencer, S. Meinert, T. Hahn, U. Dannlowski, D. Grotegerd, M. Ciccarelli, F. Iasevoli, G. Pontillo, G. D. Pearlson, D. Cobia, F. Piras, N. Banaj, D. Vecchio, M. E. A. Barendse, N. E. M. van Haren, H. J. Jo, K. Sim, Y. Quidé, M. J. Green, R. Slate, G. Cecere, W. Omlor, S. Homan, P. Homan, S. I. Thomopoulos, A. Manzari, A. Bellomo, J. A. Turner, T. G. M. van Erp, P. M. Thompson, A. Bertolino, G. Pergola, Thalamocortical Structural Covariation Networks Are Related to Familial Risk for Schizophrenia in the Context of Lower Nuclei Volume Estimates in Patients: An ENIGMA Study. Biological Psychiatry 98, 698–711 (2025).

59. N. F. Forbes, L. A. Carrick, A. M. McIntosh, S. M. Lawrie, Working memory in schizophrenia: a meta-analysis. Psychol Med 39, 889–905 (2009).

60. T. Grent-’t-Jong, J. Gross, J. Goense, M. Wibral, R. Gajwani, A. I. Gumley, S. M. Lawrie, M. Schwannauer, F. Schultze-Lutter, T. Navarro Schröder, D. Koethe, F. M. Leweke, W. Singer, P. J. Uhlhaas, Resting-state gamma-band power alterations in schizophrenia reveal E/I-balance abnormalities across illness-stages. eLife 7, e37799.

61. M. J. Minzenberg, A. J. Firl, J. H. Yoon, G. C. Gomes, C. Reinking, C. S. Carter, Gamma Oscillatory Power is Impaired During Cognitive Control Independent of Medication Status in First-Episode Schizophrenia. Neuropsychopharmacology 35, 2590–2599 (2010).

62. D. Doischer, J. Aurel Hosp, Y. Yanagawa, K. Obata, P. Jonas, I. Vida, M. Bartos, Postnatal Differentiation of Basket Cells from Slow to Fast Signaling Devices. J Neurosci 28, 12956–12968 (2008).

63. T. Miyamae, K. Chen, D. A. Lewis, G. Gonzalez-Burgos, Distinct Physiological Maturation of Parvalbumin-Positive Neuron Subtypes in Mouse Prefrontal Cortex. J. Neurosci. 37, 4883–4902 (2017).

64. A. P. Caccavano, A. Vlachos, N. McLean, S. Kimmel, J. H. Kim, G. Vargish, V. Mahadevan, L. Hewitt, A. M. Rossi, I. Spineux, S. J. Wu, E. Furlanis, M. Dai, B. L. Garcia, Y. Wang, R. Chittajallu, E. London, X. Yuan, S. Hunt, D. Abebe, M. A. G. Eldridge, A. C. Cummins, B. E. Hines, A. Plotnikova, A. Mohanty, B. B. Averbeck, K. A. Zaghloul, J. Dimidschstein, G. Fishell, K. A. Pelkey, C. J. McBain, Divergent opioid-mediated suppression of inhibition between hippocampus and neocortex across species and development. Neuron 113, 1805–1822.e7 (2025).

65. S. J. Dienel, D. A. Lewis, Alterations in cortical interneurons and cognitive function in schizophrenia. Neurobiology of Disease 131, 104208 (2019).

66. S. J. Dienel, K. E. Schoonover, D. A. Lewis, Cognitive Dysfunction and Prefrontal Cortical Circuit Alterations in Schizophrenia: Developmental Trajectories. Biol Psychiatry 92, 450–459 (2022).

67. S. M. Sherman, R. W. Guillery, Distinct functions for direct and transthalamic corticocortical connections. Journal of Neurophysiology 106, 1068–1077 (2011).

68. S. M. Sherman, Thalamus plays a central role in ongoing cortical functioning. Nat Neurosci 19, 533–541 (2016).

69. Z. Molnár, K. Y. Kwan, Development and Evolution of Thalamocortical Connectivity. Cold Spring Harb Perspect Biol 16, a041503 (2024).

70. A. C. Roberts, D. L. Tomic, C. H. Parkinson, T. A. Roeling, D. J. Cutter, T. W. Robbins, B. J. Everitt, Forebrain connectivity of the prefrontal cortex in the marmoset monkey (Callithrix jacchus): An anterograde and retrograde tract-tracing study. Journal of Comparative Neurology 502, 86–112 (2007).

71. A. S. Mitchell, S. Chakraborty, What does the mediodorsal thalamus do? Front Syst Neurosci 7, 37 (2013).

72. A. Mukherjee, N. H. Lam, R. D. Wimmer, M. M. Halassa, Thalamic circuits for independent control of prefrontal signal and noise. Nature 600, 100–104 (2021).

73. B. R. Ferguson, W.-J. Gao, Thalamic control of cognition and social behavior via regulation of GABAergic signaling and E/I balance in the medial prefrontal cortex. Biol Psychiatry 83, 657–669 (2018).

74. I.-C. Mundinano, W. C. Kwan, J. A. Bourne, Mapping the mosaic sequence of primate visual cortical development. Front Neuroanat 9, 132 (2015).

75. S. Park, K. V. Haak, S. Oldham, H. Cho, K. Byeon, B. Park, P. Thomson, H. Chen, W. Gao, T. Xu, S. Valk, M. P. Milham, B. Bernhardt, A. Di Martino, S.-J. Hong, A shifting role of thalamocortical connectivity in the emergence of cortical functional organization. Nat Neurosci 27, 1609–1619 (2024).

76. A. John, A. Saberi, A. Manoli, J. Royer, D. Y. Eriguec, V. J. Sydnor, B. Wan, S. B. Eickhoff, B. C. Bernhardt, A. Anwander, S. L. Valk, Nucleus-level thalamic organization anchors multimodal signatures of thalamocortical maturation. bioRxiv [Preprint] (2026). 10.64898/2026.07.03.736332.

77. I.-C. Mundinano, P. A. Flecknell, J. A. Bourne, MRI-guided stereotaxic brain surgery in the infant and adult common marmoset. Nat Protoc 11, 1299–1308 (2016).

78. G. Paxinos, C. R. R. Watson, M. Petrides, M. G. Rosa, H. Tokuno, The Marmoset Brain in Stereotaxic Coordinates (Elsevier, 2012; https://research.monash.edu/en/publications/the-marmoset-brain-in-stereotaxic-coordinates/).

79. J.-D. Tournier, R. Smith, D. Raffelt, R. Tabbara, T. Dhollander, M. Pietsch, D. Christiaens, B. Jeurissen, C.-H. Yeh, A. Connelly, *MRtrix3*: A fast, flexible and open software framework for medical image processing and visualisation. NeuroImage 202, 116137 (2019).

80. M. Jenkinson, C. F. Beckmann, T. E. J. Behrens, M. W. Woolrich, S. M. Smith, FSL. NeuroImage 62, 782–790 (2012).

81. B. B. Avants, N. J. Tustison, G. Song, P. A. Cook, A. Klein, J. C. Gee, A Reproducible Evaluation of ANTs Similarity Metric Performance in Brain Image Registration. Neuroimage 54, 2033–2044 (2011).

82. C. Liu, F. Q. Ye, J. D. Newman, D. Szczupak, X. Tian, C. C.-C. Yen, P. Majka, D. Glen, M. G. P. Rosa, D. A. Leopold, A. C. Silva, A resource for the detailed 3D mapping of white matter pathways in the marmoset brain. Nat Neurosci 23, 271–280 (2020).

83. R. Pomponio, G. Erus, M. Habes, J. Doshi, D. Srinivasan, E. Mamourian, V. Bashyam, I. M. Nasrallah, T. D. Satterthwaite, Y. Fan, L. J. Launer, C. L. Masters, P. Maruff, C. Zhuo, H. Völzke, S. C. Johnson, J. Fripp, N. Koutsouleris, D. H. Wolf, R. Gur, R. Gur, J. Morris, M. S. Albert, H. J. Grabe, S. M. Resnick, R. N. Bryan, D. A. Wolk, R. T. Shinohara, H. Shou, C. Davatzikos, Harmonization of large MRI datasets for the analysis of brain imaging patterns throughout the lifespan. NeuroImage 208, 116450 (2020).

84. J. Schindelin, I. Arganda-Carreras, E. Frise, V. Kaynig, M. Longair, T. Pietzsch, S. Preibisch, C. Rueden, S. Saalfeld, B. Schmid, J.-Y. Tinevez, D. J. White, V. Hartenstein, K. Eliceiri, P. Tomancak, A. Cardona, Fiji: an open-source platform for biological-image analysis. Nat Methods 9, 676–682 (2012).

85. N. Atapour, K. H. Worthy, L. L. Lui, H.-H. Yu, M. G. P. Rosa, Neuronal degeneration in the dorsal lateral geniculate nucleus following lesions of primary visual cortex: comparison of young adult and geriatric marmoset monkeys. Brain Struct Funct 222, 3283–3293 (2017).

86. R. Guirado, H. Carceller, E. Castillo-Gómez, E. Castrén, J. Nacher, Automated analysis of images for molecular quantification in immunohistochemistry. Heliyon 4, e00669 (2018).

87. R. W. Cox, AFNI: software for analysis and visualization of functional magnetic resonance neuroimages. Comput Biomed Res 29, 162–173 (1996).

88. C. Liu, C. C.-C. Yen, D. Szczupak, X. Tian, D. Glen, A. C. Silva, Marmoset Brain Mapping V3: Population multi-modal standard volumetric and surface-based templates. Neuroimage 226, 117620 (2021).

89. A. Takemoto, M. Miwa, R. Koba, C. Yamaguchi, H. Suzuki, K. Nakamura, Individual variability in visual discrimination and reversal learning performance in common marmosets. Neuroscience Research 93, 136–143 (2015).

90. A. Sadoun, M. Rosito, C. Fonta, P. Girard, Key periods of cognitive decline in a nonhuman primate model of cognitive aging, the common marmoset (*Callithrix jacchus*). Neurobiology of Aging 74, 1–14 (2019).

91. A. Gramfort, M. Luessi, E. Larson, D. A. Engemann, D. Strohmeier, C. Brodbeck, R. Goj, M. Jas, T. Brooks, L. Parkkonen, M. Hämäläinen, MEG and EEG data analysis with MNE-Python. Front. Neurosci. 7 (2013).

92. P. Virtanen, R. Gommers, T. E. Oliphant, M. Haberland, T. Reddy, D. Cournapeau, E. Burovski, P. Peterson, W. Weckesser, J. Bright, S. J. van der Walt, M. Brett, J. Wilson, K. J. Millman, N. Mayorov, A. R. J. Nelson, E. Jones, R. Kern, E. Larson, C. J. Carey, İ. Polat, Y. Feng, E. W. Moore, J. VanderPlas, D. Laxalde, J. Perktold, R. Cimrman, I. Henriksen, E. A. Quintero, C. R. Harris, A. M. Archibald, A. H. Ribeiro, F. Pedregosa, P. van Mulbregt, SciPy 1.0: fundamental algorithms for scientific computing in Python. Nat Methods 17, 261–272 (2020).

